# Mitochondrial Dysfunction is a Driver for SP-2509 Drug Resistance in Ewing Sarcoma

**DOI:** 10.1101/2021.12.13.472459

**Authors:** E. John Tokarsky, Jesse C. Crow, Lillian M. Guenther, John Sherman, Cenny Taslim, Gabriela Alexe, Kathleen I. Pishas, Galen Rask, Blake S. Justis, Ana Kasumova, Kimberly Stegmaier, Stephen L. Lessnick, Emily R. Theisen

## Abstract

Expression of the fusion oncoprotein EWS/FLI causes Ewing sarcoma, an aggressive pediatric tumor characterized by widespread epigenetic deregulation. These epigenetic changes are targeted by novel lysine specific demethylase-1 (LSD1) inhibitors, which are currently in early phase clinical trials. Single agent targeted therapy often induces resistance, and successful clinical development requires knowledge of resistance mechanisms, enabling the design of effective combination strategies. Here, we used a genome-scale CRISPR-Cas9 loss-of-function screen to identify genes whose knockout (KO) conferred resistance to the LSD1 inhibitor SP- 2509 in Ewing sarcoma cell lines. Multiple genes required for mitochondrial electron transport chain (ETC) complexes III and IV function were hits in our screen. We validated this finding using genetic and chemical approaches including CRISPR KO, ETC inhibitors, and mitochondrial depletion. Further global transcriptional profiling revealed that altered complex III/IV function disrupted the oncogenic program mediated by EWS/FLI and LSD1 and blunted the transcriptomic response to SP-2509. These findings demonstrate that mitochondrial dysfunction modulates SP-2509 efficacy and suggest that new therapeutic strategies combining LSD1 with agents which prevent mitochondrial dysfunction may benefit patients with this aggressive malignancy.

## Introduction

Ewing sarcoma is the second most common bone malignancy in pediatric, adolescent, and young adult patients (Burningham, Hashibe et al., 2012, Delattre, Zucman et al., 1992). Five-year survival rates for patients with localized disease is 70-80%, but due to its highly aggressive nature, patients with metastatic, recurrent, or refractory tumors face poor outcomes with only 10- 30% long-term survival (Gaspar, Hawkins et al., 2015, Grunewald, Cidre-Aranaz et al., 2018).

Ewing sarcoma is histologically classified as a small, round, blue cell tumor, and has one of the lowest mutational burdens of any cancer (0.15 mutations/Mb) (Brohl, Solomon et al., 2014, Crompton, Stewart et al., 2014, Lawrence, Stojanov et al., 2013). Ewing sarcoma is driven by a single fusion oncogene, most commonly EWS/FLI (Delattre et al., 1992). EWS/FLI is the direct result of a *t*(11;22)(q24; q12) chromosomal translocation between the FET gene family member, Ewing sarcoma breakpoint region 1 protein (*EWSR1*), and the ETS transcription factor, Friend leukemia integration 1 transcription factor (*FLI1*). The resulting chimeric protein is a potent master transcriptional regulator and primary driver of oncogenesis (Turc-Carel, Aurias et al., 1988, Turc-Carel, Philip et al., 1984). EWS/FLI and other FET/ETS oncoproteins bind to DNA motifs containing either a consensus ETS motif (5′-*ACC<u>GGAA</u>GTG*-3′) (Szymczyna & Arrowsmith, 2000, Wei, Badis et al., 2010) or microsatellites consisting of GGAA repeats (Johnson, Mahler et al., 2017). Binding to these DNA motifs allows EWS/FLI to promote chromatin accessibility and establish *de novo* enhancers (Riggi, Knoechel et al., 2014). In addition, the low complexity domain of EWSR1 in EWS/FLI recruits chromatin regulators, such as BAF, NuRD, and p300 which are important for EWS/FLI-mediated activation and repression (Boulay, Sandoval et al., 2017), (Sankar, Bell et al., 2013).

Lysine-specific demethylase-1 (LSD1) is a flavin adenine dinucleotide (FAD)-dependent amine oxidase that is highly expressed in Ewing sarcoma (Bennani-Baiti, Machado et al., 2012, Pishas, Drenberg et al., 2018, Shi, Lan et al., 2004). LSD1 catalyzes the demethylation of mono- and di- methyl lysines at H3K4 and H3K9 (Shi et al., 2004) and is recruited by EWS/FLI as a member of the NuRD (Wang, Zhang et al., 2009), and REST corepressor (CoREST) (Chen, Yang et al., 2006) complexes. We have previously demonstrated that targeting LSD1 with the small molecule inhibitor SP-2509 is efficacious in Ewing sarcoma pre-clinical models (Sankar, Theisen et al., 2014, Sorna, Theisen et al., 2013, Theisen, Pishas et al., 2016). SP-2509 is a reversible, allosteric inhibitor of LSD1 that halts Ewing sarcoma cell proliferation, induces apoptosis, and blocks tumor growth in pre-clinical Ewing sarcoma models (Pishas et al., 2018, Sankar et al., 2013, Sankar et al., 2014). At present, an analog of SP-2509, known as seclidemstat (SP-2577, *Salarius Pharmaceuticals*), is currently in clinical trials to treat relapsed or refractory Ewing sarcoma (NCT03600649).

Acquired resistance to single agent small molecules is a challenge in the development of novel cancer therapies. Given that small molecules targeting LSD1 could have an impact in patients with this aggressive disease, developing combination strategies to aid future clinical trials requires a deeper understanding of potential drug resistance pathways. Our laboratory previously generated an SP-2509 resistant Ewing sarcoma cell line from parental A673 cells through chronic exposure to escalating concentrations of SP-2509 (Pishas & Lessnick, 2018). This resistant A673 cell line demonstrated a 55-fold greater IC50 over parental A673 cells (7.5 µM vs. 0.18 µM, respectively). Following removal of SP-2509 treatment, a ten-fold increased IC50 (2 µM) was maintained upon re-challenge, suggesting a non-reversible phenotype. No LSD1 mutations were identified in these cells, and instead a mutation in mitochondrial ribosomal protein L45 (*MRPL45*) was found in 85% of the drug resistant cell population.

An emerging use for CRISPR screening technology is to determine genes whose deletion causes resistance to a drug of interest, thereby suggesting novel mechanisms involved in drug activity and potential escape pathways in cancer cells (Sulahian, Kwon et al., 2019). We undertook a genome-scale CRISPR-Cas9 loss-of-function screen in two Ewing sarcoma cell lines in the presence of SP-2509. This identified genes involved in drug resistance related to mitochondrial function, specifically in the electron transport chain (ETC) complexes III (CIII) and IV (CIV).

This was subsequently validated utilizing genetic KO models, ETC inhibitors, and mitochondrial depletion. SP-2509 treatment causes global changes in the transcriptome of Ewing sarcoma cells, and transcriptomic investigation of resistant cells using RNA-sequencing (RNA-seq) revealed that resistant cells show a blunted transcriptional response. We further report that the mitochondrial dysfunction that cause SP-2509 drug resistance also alters the transcriptional program enforced by EWS/FLI and LSD1. We propose that combining LSD1 inhibitors with agents that promote oxidative phosphorylation may be useful to circumvent these changes and prevent development of acquired resistance in Ewing sarcoma models.

## Results

### A genome-scale CRISPR-Cas9 screen identifies unique mitochondrial gene deletions conferring SP-2509 resistance in Ewing sarcoma cells

Given the potential of LSD1 inhibitors for treating patients with Ewing sarcoma, we employed an unbiased genomic screening approach to identify mechanisms of resistance to SP-2509. We performed a genome-scale CRISPR-Cas9 loss-of-function screen (Figure 1A) using the Avana-4 lentiviral library (Doench, Fusi et al., 2016, Meyers, Bryan et al., 2017) which targets 18,333 human genes, with 74,378 sgRNAs, each carrying a unique barcode (Supplementary File 1).

**Figure 1.**
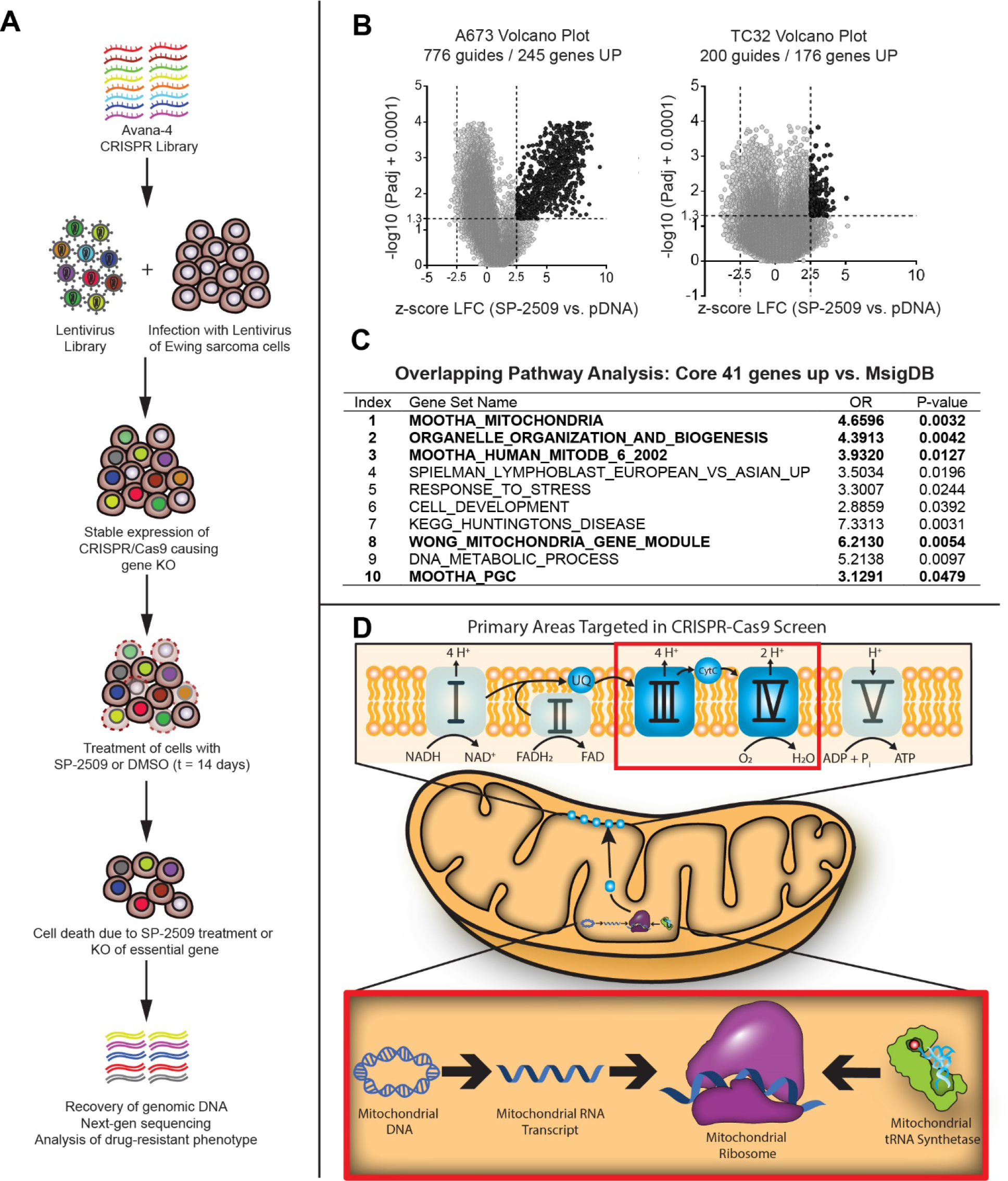
CRISPR-Cas9 Screen Identifies Mitochondrial Dysfunction as a Drug-Resistance Phenotype. **A.** Overview of CRISPR-Cas9 screen. An Avana-4 library was used to generate a lentivirus library that was used to infect A673 and TC-32 cells that stably expressed Cas9. After antibiotic selection, cells were treated with either DMSO or 300 nM SP-2509 for 14 days. Cells remaining after 14 days were collected, and their DNA was purified. PCR and barcoding were performed, followed by next-generation sequencing to identify the genes whose KO allowed cells to survive SP-2509 treatment. **B.** Volcano plots for A673 and TC-32 cells. Gene hits that had a log2Fold (LFC) z-score > 2.5, and an -log10 p-adjusted value of 0.05 (black dots) were considered significant. **C.** Overlapping analysis of hits from A673 and TC-32, identified 41 core gene hits whose KO promoted resistance to SP-2509. These 41 genes were used to identify genesets from MsigDB, that were common between both cell lines. **D.** Representative image of the categories of mitochondrial genes that were identified in our CRISPR-Cas9 screen, including mitochondrial ribosomal proteins, mitochondrial tRNA synthetases, mitochondrial DNA-replication and transcription machinery, and ETC complexes III and IV. These components are surrounded by red boxes.

After transducing two SP-2509 sensitive cell lines, A673 and TC-32 (Pishas et al., 2018), cells were treated with SP-2509 (300 nM) or equivalent DMSO control, with the concentration optimized to promote positive selection of resistant cells. For each condition, cells were harvested at both an early time point (day 0) and at 14 days. Genomic DNA was then isolated from both treatment arms and barcodes were sequenced (Figure 1A). The sgRNAs that provided a growth advantage with SP-2509 were enriched in the treated arm compared with DMSO controls. A core list of genes whose KO conferred resistance to SP-2509 was generated by examining z-scores calculated on sgRNA barcode reads by normalizing to an early time point. Using a log2 fold change (log2FC) expression of > 2.5 and a significance value (p < 0.05) cutoff, we identified 245 genes and 176 genes in A673 and TC-32 cells, respectively, whose KO conferred SP-2509 resistance (Figure 1B; Supplementary File 2).

Of these genes, 41 were common hits across both A673 and TC-32 cells (Supplementary Table S1). We analyzed these common genes for phenotypic signatures in the Molecular Signatures Database (MSigDB) which revealed mitochondrial metabolism, organelle organization and biogenesis, DNA damage/repair, and apoptosis pathways significantly enriched in the SP-2509 resistant state (*P<0.05*) (Figure 1C). Interestingly, 5 out of the top 10 most significantly enriched signatures were mitochondrial-related pathways (Figure 1C). The loss of genes encoding mitochondrial ribosomal proteins (MRPs), CIII and CIV proteins, and mitochondrial tRNA synthetases, but not mutations in ETC complexes I, II or V (CI, CII, CV), conferred SP-2509 resistance, suggesting specificity to particular pathways and not simply broad mitochondrial dysfunction (Table 1 and Figure 1D). Importantly, our CRISPR screen independently confirmed our previous finding of a stop-gain *MRPL45* mutation promoting acquired resistance to SP-2509 in A673 cells (Pishas & Lessnick, 2018). In total, 74/78 possible MRPs and 18/56 possible mitochondrial tRNA synthetases (Table 1) were identified as positive hits in our CRISPR screen. MRPs and mitochondrial tRNA synthetases are required for translation of the human mitochondrial genome, which consists of 37 total genes that encode for 13 proteins, 22 tRNAs, and 2 rRNAs (Taanman, 1999). The majority of the human mitochondrial genome encodes for proteins that form core subunits of ETC complexes I, III, IV, and V (Gammage & Frezza, 2019). Other nuclear-encoded genes that are important for replication, transcription, and translation of the mitochondrial genome were also found in our CRISPR screen (Supplementary File 2), including *POLG/G2* (polymerase gamma), and *POLRMT* (mitochondrial RNA polymerase).

**Table 1:**
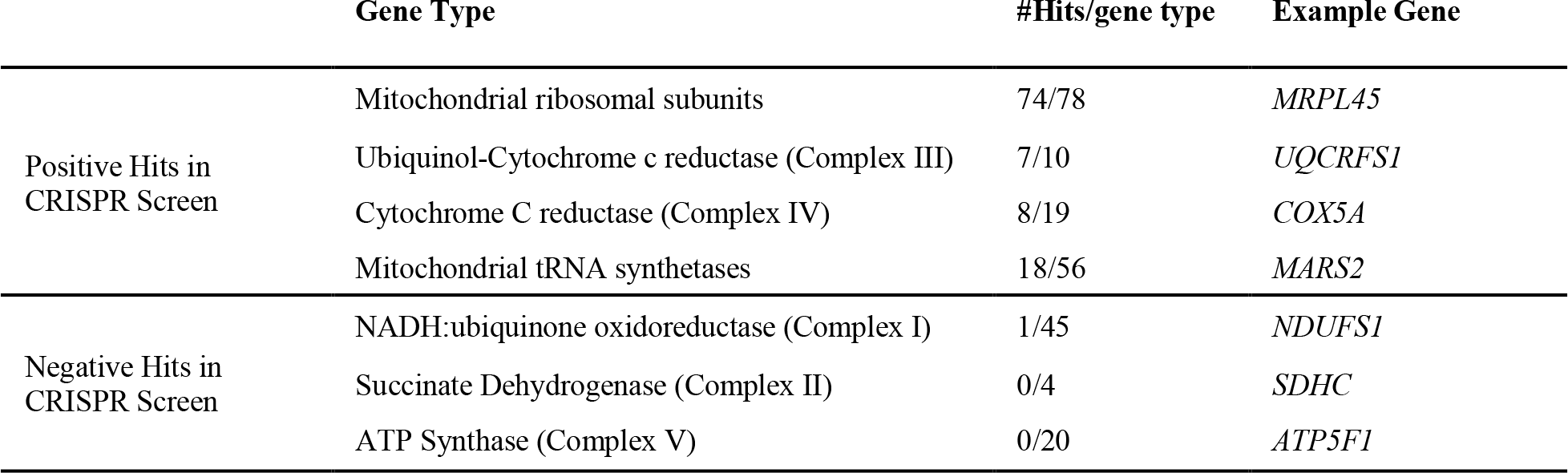
CRISPR-Cas9 screen gene hits clustered based on mitochondrial gene groups.

To validate this CRISPR screen, we generated KO of various genes of interest from the mitochondrial ribosome – *MRPL45*; CIII – Ubiquinol-Cytochrome C Reductase, Rieske Iron-Sulfur Polypeptide 1 (*UQCRFS1*) and cytochrome C1 (*CYC1*); and an CIV assembly factor – Cytochrome C Oxidase Assembly Factor 4 Homolog (*COA4*). We also generated KOs for CI – NADH:Ubiquinone Oxidoreductase Core Subunit S1 (*NDUFS1*), CII – Succinate Dehydrogenase C (*SDHC*), and CV – ATP synthase F(0) complex subunit B1 (*ATP5F1*). Cell viability assays were performed to test the efficacy of SP-2509 on these KO cells. We used sgRNA sequences identical to those found in the Avana-4 CRISPR library, and KO was confirmed by western blot (Supplementary Figure S1). Mitochondrial hits from the CRISPR screen, including *MRPL45*, *UQCRFS1*, *CYC1,* and *COA4* either significantly increased the SP- 2509 IC50 or trended toward an increase in polyclonal KO populations (Supplementary Figure S1). In contrast, polyclonal populations with sgRNAs targeting ETC components from CI (*NDUFS1*), CII (*SDHC*), and CV (*ATP5F1*) do not show any significant changes in SP-2509 IC50 (Supplementary Figure S1). This preliminary validation effort confirmed the specificity of our CRISPR screen to particular components (MRPs, CIII and CIV) of the mitochondria rather than general mitochondrial dysfunction.

### Single cell clones exhibiting KO of mitochondrial components show enhanced resistance to SP- 2509

We found it notable that the specificity of hits enriching for CIII and CIV of the ETC, and the convergence with our prior studies (Pishas & Lessnick, 2018), suggests that mitochondrial dysfunction is a bona fide driver of SP-2509 resistance. To better define the link between mitochondrial dysfunction and resistance to SP-2509, we isolated single cells following lentiviral transduction with CRISPR constructs, grew out monoclonal populations, and screened for clones containing complete gene loss (Figure 2A). This method generated monoclonal cell populations containing KO of the following genes: *MRPL45* (MRPL45 KO); *UQCRFS1* (UQCRFS1 KO) and *CYC1* (CYC1 KO).

**Figure 2.**
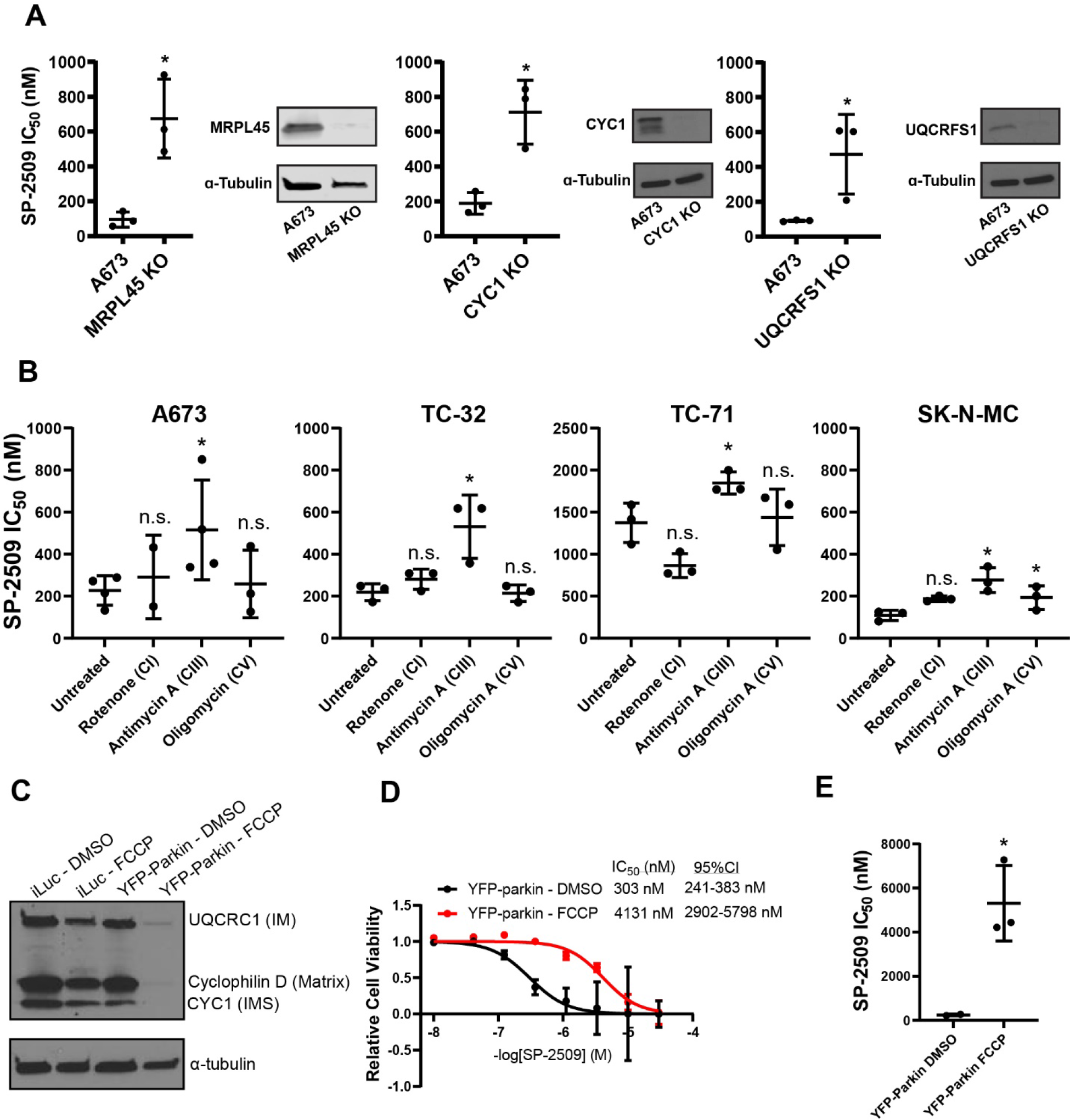
Validation of CRISPR screen through genetic KO, chemical inhibition, and mitochondrial depletion. **A.** Compiled SP-2509 IC50 data and western blot images for monoclonal KO cells; mitochondrial ribosomal protein L45 (MRPL45 KO), Ubiquinone-Cytochrome C Reductase, Rieske Iron- Sulfur Polypeptide 1 (UQCRFS1 KO), Cytochrome C1 (CYC1 KO). Compiled data is from n=3 independent experiments. Statistical analysis was performed by using a student’s t-test. Data that reached a significance (p-value) of ≤ 0.05 are denoted with *. **B.** Compiled SP-2509 IC50 data for simultaneous treatment of indicated cell lines with SP-2509 and ETC inhibitors, rotenone (CI), antimycin A (CIII), or oligomycin (CV). Statistical analysis was performed by using a student’s t-test comparing each inhibitor treatment with the untreated cell lines, with * denoting a significance (p-value) of ≤ 0.05, and values that are not significant denoted as n.s. **B.** Western blot analysis of mtø cells for mitochondrial proteins using Membrane Integrity WB Antibody Cocktail (abcam; ab110414). Samples are denoted with A673 cells infected with retroviral constructs for YFP-Parkin and iLuc (mock infection), that were treated with either DMSO or FCCP for 48 hr. Blots for Ubiquinone-cytochrome c reductase core protein 1 (UQCRC1) represents the mitochondrial intermembrane (IM), Cyclophilin D in the mitochondrial matrix, cytochrome C1 (CYC1) in the mitochondrial intermembrane space (IMS), and α-tubulin was used as a loading control. Depletion of these proteins in the YFP-Parkin – FCCP condition indicates depletion of mitochondria from the cells. **C.** Dose response curve comparing A673 cells expressing YFP-parkin either treated with DMSO or FCCP. **D.** Compiled SP-2509 IC50 data for mtø cells. Statistical analysis was performed by using a student’s t-test comparing each inhibitor treatment with the untreated cell lines, with * denoting a significance (p-value) of ≤ 0.05.

We first analyzed the impact of these gene KOs on mitochondrial function in our monoclonal populations using a seahorse instrument. This determined that both MRPL45 KO and CYC1 KO had significantly decreased basal respiration, whereas UQCRFS1 KO cells retained similar basal and maximal respiration to parental A673s (Supplementary Figure S2). Cell viability assays indicated that each monoclonal KO cell line showed enhanced resistance to SP-2509 (Figure 2A). This further confirmed the findings from the CRISPR screen and highlighted the ability of significantly reduced mitochondrial gene activity to dampen response to SP-2509.

### Electron transport chain inhibitors targeting CIII, but not CI or CV, confer SP-2509 resistance in Ewing sarcoma cell lines

Having validated the CRISPR screen using genetic approaches, we next assayed whether loss of specific ETC complexes led to SP-2509 resistance using chemical probes targeting different complexes in the ETC. We used rotenone to block CI, antimycin A to block CIII, and oligomycin A to block CV. Initial optimization experiments identified 50 nM rotenone, 500 nM antimycin A, or 500 nM oligomycin A as treatment conditions that could achieve sufficient ETC blockade during a 72-hr treatment (Supplementary Figure S3). We then performed cell viability assays under dual treatment with SP-2509 (dose range 0.01 – 30 µM) and the various ETC inhibitors. We found that only antimycin A conferred drug resistance to SP-2509 compared to untreated cells in all 4 Ewing sarcoma cell lines tested (A673, TC-32, SK-N-MC, and TC-71 (Figure 2B). In contrast, neither CI nor CV inhibition significantly or consistently altered SP- 2509 IC50 values (Figure 2B). These results lend additional validation to the CRISPR screen findings, with chemical inhibition of the ETC at CIII, but not CI or CV, inducing resistance to SP-2509 in Ewing sarcoma cells.

### Loss of mitochondria leads to resistance to SP-2509 in A673 cells

Genetic and chemical approaches validated our initial findings that specific components of the mitochondria, including mitochondrial transcriptional and translational machinery and ETC complexes CIII and CIV, are important for sensitivity to SP-2509, and loss of these components confers resistance. We therefore asked whether resistance arose following the loss of a specific mitochondrial function, or instead as a result of residual organelle dysfunction. To address this question, we depleted mitochondria from cells using mitophagy, generating mitochondrial null (mtø) A673 cells. In this system, YFP-tagged Parkin (YFP-Parkin), an E3 ubiquitin ligase, is retrovirally transduced and mitochondrial stress is caused with the protonophore FCCP. YFP-Parkin localizes to the stressed mitochondria and induces mitophagy (Supplementary Figure S4A) (Correia-Melo, Ichim et al., 2017). We confirmed mitochondrial depletion via western blot with a mitochondrial antibody cocktail recognizing proteins in the inner-membrane (IM), the inner-membrane space (IMS), and the mitochondrial matrix (Figure 2C). Retroviral transduction with a mock control (iLuc) confirmed that FCCP alone did not deplete mitochondria (Figure 2C) or promote resistance to SP-2509 (Supplementary Figure S4B). We performed a Seahorse assay to confirm mtø cells had lost their oxidative capacity (Supplementary Figure S4C). Cell viability assays performed after SP-2509 treatment in our mtø cells showed a mean IC50 of 5309 nM, 21- fold higher than the mean IC50 observed with control cells (mean IC50 = 244 nM; p-value = 0.02) (Figure 2D-E). These data support that the loss of specific mitochondrial functions, rather than the resulting cellular dysfunction, contributes to SP-2509 resistance.

### RNA-sequencing reveals KO clones show differing response to SP-2509

We have previously shown that LSD1 enforces EWS/FLI-mediated transcription and that SP- 2509 treatment reverses EWS/FLI-mediated gene regulation (Pishas et al., 2018), (Theisen, Selich-Anderson et al., 2021). Given that the functional relevance of mitochondrial status on LSD1 and EWS/FLI activity is unknown, and that mitochondrial dysfunction imparted resistance to SP-2509, we next tested whether resistant cells showed altered LSD1-EWS/FLI activity and how the transcriptional response to SP-2509 in resistant cells differed from sensitive cells.

Following 48 hrs of treatment with either vehicle or 500 nM SP-2509, parental A673 cells, MRPL45 KO cells, CYC1 KO cells, and UQCRFS1 KO cells were assayed by RNA-sequencing (RNA-seq) and differentially expressed genes (DEGs) were determined (Figure 3A). As has been shown previously in A673 cells, SP-2509 treatment altered the expression of thousands of genes (12125 total DEGs; 6929 upregulated, 5196 downregulated). Interestingly, we saw differing transcriptional responses among our monoclonal cell lines. UQCRFS1 KO cells showed the greatest number of DEGs (11528 total DEGs; 6373 upregulated, 5155 downregulated), comparable to A673. Both CYC1 KO cells (7351 total DEGs; 3855 upregulated, 3496 downregulated) and MRPL45 KO cells showed fewer DEGs (4681 DEGs total; 2925 upregulated, 1756 downregulated) suggesting a blunted transcriptional response. Principal component analysis (PCA) revealed that vehicle-treated CYC1 KO, UQCRFS1 KO, and A673 cells cluster together, while vehicle-treated MRPL45 KO cells cluster apart from this group along the first principal component (PC1) axis (Figure 3B). SP-2509 treatment generally caused a shift along the principal component 2 (PC2) axis, with variable effects on each cell type. The largest shift was seen for parental A673 cells and UQCRFS1 KO cells, with lesser effects on CYC1 KO cells and an even further diminished response for MRPL45 KO cells. These effects agree with the total number of DEGs observed in each cell line and suggest a spectrum of transcriptional response, with the most resistant cells showing the smallest change in transcriptome, and vice versa.

**Figure 3.**
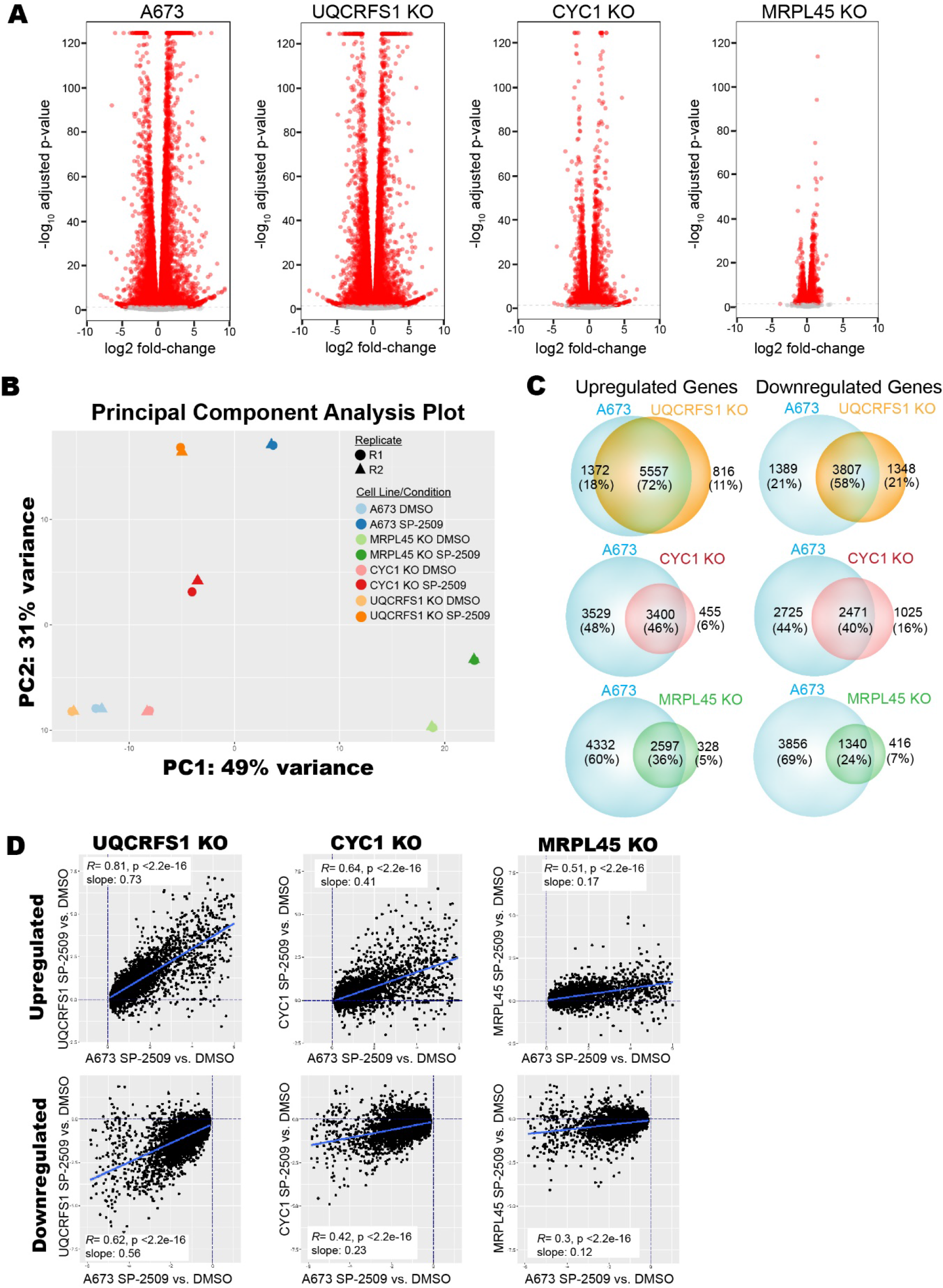
RNA-sequencing of monoclonal mitochondrial KO cell lines. **A.** Volcano plots comparing SP-2509 treatment vs. DMSO treatment for each indicated cell line. Each dot represents a differentially expressed gene (DEG) with significantly changed DEGs (red) having a -log10 adjusted p-value < 0.05, and non-significant DEGs (grey) failing to reach significance. **B.** Principal component analysis (PCA) plot of the transcriptional profiles of indicated monoclonal KO cell lines and parental A673 cells. Principal component 2 (PC2) on the y-axis is plotted against principal component 1 (PC1) on the x-axis. Different cell conditions are depicted by color and replicates are represented with different shapes. **C.** Venn diagrams comparing the overlap of differentially expressed genes caused by treatment with SP-2509 for A673 vs. the indicated monoclonal KO cell line. **D.** Scatterplots depicting common genes differentially regulated during treatment with SP-2509 for parental A673 cells versus each indicated monoclonal KO cell line. Data is plotted as log2FC of each monoclonal KO cell on the y-axis against the log2FC of A673 parental cells on the x-axis. Dotted lines depict x = 0 and y = 0. Significance was defined as Benjamini-Hochberg adjusted p < 0.05 with no fold-change cutoff. Lines of best fit were derived from the linear model only for genes with significant changes in both parental and monoclonal KO cells to represent the common DEGs. Pearson correlation coefficients and their p-values are depicted with the slope from the linear model in the boxed inset.

Because we observed that all cell lines shifted in the same direction along PC2 to varying degrees following SP-2509 treatment, we next hypothesized that similar genes were changing in response to treatment. To address this, we used Venn analysis to compare the SP-2509-induced DEGs in A673 cells to those identified in the various monoclonal KO cell lines (Figure 3C). We found that a majority of the DEGs in each KO cell line were shared with A673 cells (Figure 3C). We noted that the more resistant cell lines had a smaller number of DEGs and ahigh degree of overlap between DEGs in KO lines and A673s. Therefore, we next asked whether decreased responsiveness arose because either a subset of genes failed to respond to treatment or that all genes decreased in their responsiveness resulting in a lower number of genes exceeding the cutoffs to define a DEG. To do this, we plotted the observed fold change for a given gene in a monoclonal KO cell line against the fold change observed for that gene in A673 cells, separating up- and downregulated genes (Figure 3D). Significant correlation was observed between the transcriptional response in all KO cell lines with that in A673 cells (all p-values < 2.2x10^-16^), while the strength of the correlation, measured by both the *R* value and the slope of the line of best fit, varied across cell lines. As suggested by the number of DEGs, PCA analysis, and the Venn analysis, UQCRFS1 KO showed the strongest correlation in both upregulated and down regulated genes (*R*=0.81 slope =0.83, and *R*=0.62 slope=0.56, respectively). This was then followed by CYC1 KO (upregulated: *R*=0.64, slope =0.41; downregulated: *R*=0.42, slope=0.23), while MRPL45 KO showed the weakest correlation (upregulated: *R*=0.51, slope =0.17; downregulated: *R*=0.30, slope=0.12). This demonstrates that as you move from least resistant (UQCRFS1 KO) to more resistant (CYC1 KO) to most resistant (MRPL45 KO) the transcriptional response to SP-2509 is globally diminished, akin to turning down the “volume” of this response. Together, these data show that though SP-2509 largely regulates the same genes in A673 cells and mitochondrial KO lines, altering the mitochondrial function in A673s blunts the response to SP-2509.

### Mitochondrial dysfunction induces transcriptional changes that mimic SP-2509 treatment

We reasoned that the observed decrease in transcriptional response to SP-2509 could arise either because the genes regulated by SP-2509 were more resistant to change or because KO of mitochondrial genes had already altered expression of these genes. To further distinguish between these possibilities, we first identified the basal transcriptional changes induced by KO in all three monoclonal cell lines. This revealed that MRPL45 KO had the greatest DEGs with respect to A637 cells (12098 total DEGs; 6163 upregulated, 5935 downregulated), followed by CYC1 KO (8026 total DEGs; 4632 upregulated, 3384 downregulated), and then UQCRFS1 KO (6868 total DEGs; 3434 upregulated, 3434 downregulated), in agreement with the PCA in Figure 3B (Supplementary Table S2). Interestingly, the number of DEGs observed in the most resistant MRPL45 KO cells was comparable to that observed for SP-2509-treated A673 cells. We then used Gene Set Enrichment Analysis (GSEA) (Subramanian, Tamayo et al., 2005) to investigate whether the DEGs caused by mitochondrial dysfunction were functionally related to those DEGs caused by SP-2509 treatment in A673 cells. Using |normalized enrichment score (NES)| > 1.5 as a cutoff for significance, GSEA revealed that upregulated and downregulated genes in MRPL45 KO cells (NES: 2.9 and -3.6, respectively) shared significant functional similarity with SP-2509 treatment in A673 cells (Figure 4A). Both UQCRFS1 KO (upregulated NES: 1.5; downregulated NES: -2.9) and CYC1 KO (upregulated NES: 1.7; downregulated NES: -2.9) displayed functional similarity to SP-2509 treatment, albeit with lower NESs when upregulated genes were analyzed (Figure 4B and Supplementary Figure S5). These data indicate that our monoclonal KO cell lines have undergone significant transcriptional changes that resemble treatment with SP- 2509, supporting a model wherein the transcriptional response to SP-2509 is diminished in resistant cells because the transcriptome already resembles that of a treated cell.

**Figure 4.**
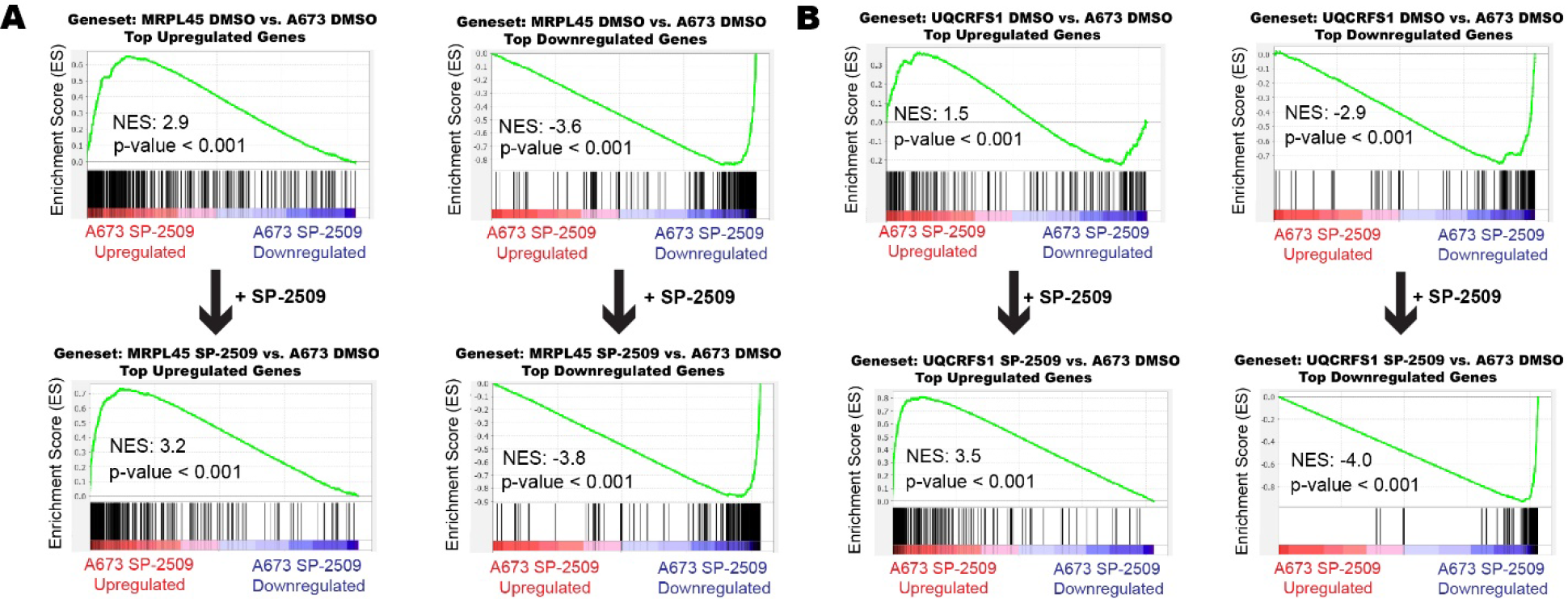
Gene Set Enrichment Analysis (GSEA) of monoclonal KO cell lines compared to SP-2509 treated A673 cells. Gene set enrichment analysis (GSEA) from RNA-seq experiments using differentially expressed genes (DEGs) for A673 cells treated with SP-2509 (A673 SP-2509 vs. A673 DMSO) as the rank-ordered list (ROL). The ROL was used for each comparison to a geneset for MRPL45 KO (**A**) and UQCRFS1 KO (**B**) of the top upregulated and downregulated genes. The top panel represents comparisons for the 480 most upregulated and downregulated genes for the indicated monoclonal KO treated with DMSO vs. A673 DMSO. The bottom panels represents GSEA for the most upregulated and downregulated genes for the indicated monoclonal KO treated with SP- 2509 vs. A673 DMSO. Arrows indicate changes in gene expression from DMSO treated cells to SP-2509 treated cells. Normalized enrichment scores (NES) and p-values are shown for each GSEA.

Taken together, our transcriptomic data suggest that our most resistant MRPL45 KO cells displayed a basal transcriptome that most resembled SP-2509 treatment and had the weakest transcriptional response to SP-2509 treatment, and vice versa with our least resistant UQCRFS1 KO cells. We next hypothesized that the combination of basal transcriptional changes induced by mitochondrial dysfunction along with further transcriptional changes induced by SP-2509 treatment would more completely recapitulate the SP-2509 transcriptional signature of sensitive A673 cells. To address this question, we used GSEA to compare the DEGs from monoclonal KO cells treated with SP-2509 with respect to parental A673 cells to the A673 SP-2509 transcriptional signature. Interestingly, for our MRPL45 KO cells treated with SP-2509 (Figure 4A; bottom panel), there was no substantial change in NES scores for upregulated genes (NES: 2.9 → 3.2) or downregulated genes (NES: -3.6 ◊ -3.8). This analysis supports that the MRPL45 KO basal transcriptional profile largely resembled an A673 cell treated with SP-2509 and is consistent with the observation that MRPL45 KO cells have the most blunted response to SP- 2509 treatment. In contrast, UQCRFS1 KO cells (Figure 4B; bottom panel) showed an increased functional enrichment for both upregulated (NES: 1.5 → 3.5) and downregulated (NES: -2.9 ◊ - 3.9) genes. Similar results were obtained for CYC1 KO cells (Supplementary Figure S5; upregulated genes - NES: 1.7 → 3.1; downregulated genes - NES: -2.9 ◊ -3.8). These analyses demonstrate that transcriptional regulation due to KO of mitochondrial genes resemble SP-2509 treatment to varying degrees generally correlating with the level of resistance. Less resistant cells CYC1 KO and UQCRFS1 KO cells show further transcriptional changes in response to SP-2509 treatment.

Since the transcriptome of our monoclonal KO cell lines resembled an A673 cell treated with SP-2509 and because SP-2509 reverses EWS/FLI transcriptional activity, we next hypothesized that mitochondrial dysfunction also disrupted EWS/FLI transcriptional activity (Pishas et al., 2018, Sankar et al., 2014). We used GSEA to compare the transcriptional changes associated with our mitochondrial KO lines with the EWS/FLI transcriptional signature. This revealed that EWS/FLI transcriptional activity is decreased in both MRPL45 KO (upregulated NES: -2.8; downregulated NES: 2.3), and CYC1 KO (upregulated NES: -2.2; downregulated NES: 1.9) (Supplementary Figure S6). In contrast, UQCRFS1 KO results in upregulation of a subset of EWS/FLI-repressed genes (upregulated NES: -2.0) but has limited impact in blunting EWS/FLI-mediated gene activation (downregulated NES: -2.1). Since LSD1 and EWS/FLI regulate similar targets, we expanded this analysis to LSD1-mediated gene regulation (Pishas et al., 2018).

Compared to EWS/FLI-mediated transcriptional activity, LSD1-mediated transcriptional activity was more impacted by mitochondrial dysfunction in MRPL45 KO cells (upregulated NES: -4.1; downregulated NES: 3.8) and CYC1 KO cells (upregulated NES: -3.1; downregulated NES: 2.0). Again, in contrast, UQCRFS1 KO demonstrated no reversal of the LSD1 transcriptional activity (upregulated NES: 1.9; downregulated NES: -3.7). Together, these data indicate that both EWS/FLI and LSD1 require mitochondrial activity in Ewing sarcoma cells, and that mitochondrial dysfunction disrupts EWS/FLI- and LSD1-mediated gene regulation. This disruption may explain a reduction in oncogenic potential observed in MRPL45 KO and CYC1

KO cells, but not UQCRFS1 KO cells (Supplementary Figure S7). Notably, though both EWS/FLI (Sankar et al., 2014) and LSD1 depletion confer resistance to SP-2509 treatment (Supplementary Figure S8A-C), and we show that EWS/FLI and LSD1 protein levels are stable across the KO cell lines tested here (Supplementary Figure S8D-E). This suggests alternative mechanisms may link mitochondrial function to transcriptional regulation in Ewing sarcoma cells and this is an important area for future studies.

## Discussion

Effective targeted therapies are urgently needed to treat recurrent or refractory Ewing sarcoma, and a promising clinical molecule, seclidemstat, may help these patients. Unfortunately, clinical use is often hindered by intrinsic- or acquired-drug resistance that renders targeted therapeutics ineffective (Groenendijk & Bernards, 2014, Sarmento-Ribeiro, Scorilas et al., 2019). Identifying and understanding potential drug resistance pathways will aid in the development of combinational therapies that suppress resistance, or screening approaches to identify the patients with greatest potential for benefit. As we have addressed in this report and previously, (Pishas & Lessnick, 2018), drug resistance arising through mitochondrial dysfunction may hinder efficacy in patients with Ewing sarcoma. In this study, we demonstrate through genomic screening and validate using genetic and chemical approaches, that SP-2509 drug resistance is mediated by mitochondrial dysfunction in Ewing sarcoma. We further demonstrate that mitochondrial dysfunction causes transcriptional changes in Ewing sarcoma that mimic EWS/FLI and LSD1 depletion and thus blunts the cellular response to SP-2509.

Mitochondria act as the key metabolic hub for the cell by producing ATP and other macromolecule precursors that aid in cell proliferation. Additionally, many studies have shown that mitochondria act as a key signaling hub and interact with the nucleus by generating metabolites important for chromatin regulation, a process known as retrograde signaling (Liu & Butow, 2006, Martinez-Reyes & Chandel, 2020, Yang & Kim, 2019). For example, a subset of gastrointestinal stromal tumors (GIST) is caused by mutations in succinate dehydrogenase (CII) enzymes, resulting in an accumulation of succinate which leads to inhibition of Ten-eleven translocation (TET) dioxygenases and dysregulation of histone demethylation (Killian, Kim et al., 2013, Mason & Hornick, 2013). Mitochondrial metabolites such as acetyl-CoA, succinate, fumarate, α-ketoglutarate (α-KG), S-adenosyl methionine (SAM) and the oncometabolite, L-2- HG, are all responsible for regulating chromatin modifying enzymes such as histone acetylases, histone demethylases such as Jumonji C domain-containing proteins, and DNA demethylases such as TET dioxygenases (Chakrabarty & Chandel, 2021, Martinez-Reyes & Chandel, 2020). Changes in the mitochondrial ETC proteins also can have profound effects on the function of cells. For example, regulatory T cells with CIII deficiency led mice to develop systemic inflammation, thymic atrophy, enlargement of lymph nodes, significantly activated CD4^+^ and CD8^+^ T cells, and impaired viability past 3 weeks of age (Weinberg, Singer et al., 2019). The CIII deficient regulatory T cells had an increase in DNA hypermethylation and gene expression changes via the buildup of 2-HG and succinate, and a decreased ratio of NAD^+^/NADH (Weinberg et al., 2019). Currently, it is unclear how mitochondria in Ewing sarcoma are involved in altering chromatin and transcriptional regulation through LSD1, or otherwise. In this study, we observed large changes in basal gene expression for our monoclonal KO cell lines indicating that transcriptional regulation is dramatically altered by mitochondrial dysfunction (Figure 3 and Supplementary Table S2). Furthermore, this differential regulation of gene expression in our monoclonal KO cells appears to blunt the response to SP-2509 in Ewing sarcoma cells.

Though this study examines potential routes of SP-2509-resistance in Ewing sarcoma, the mechanism by which drug resistance arises is not fully understood. In Figure 5A, we propose a preliminary model with each arrow representing a distinct (but not exclusive) situation where mitochondria could potentially influence SP-2509 treatment. First, mitochondria containing functional CIII and CIV could directly activate SP-2509, through chemical modification, and this activated SP-2509 could be required to inhibit LSD1. Secondly, as has been shown in other studies (Chakrabarty & Chandel, 2021, Diebold, Gil et al., 2019, Martinez-Reyes & Chandel, 2020, Weinberg et al., 2019), mitochondrial metabolites could play a role in changing the epigenetic marks on histones. This could potentially alter the function and/or localization of LSD1 (or other enzymes) on chromatin, thus influencing the epigenetic landscape of the cell and this potential mechanism is the subject of ongoing studies. Lastly, mitochondria may be influencing the basal gene expression of each cell, which is a phenomenon we observed with our monoclonal cells (Figure 3 and 4). Adopting a transcriptome that resembles an SP-2509 treated cell likely requires the engagement of compensatory mechanisms to prevent cell death.

**Figure 5.**
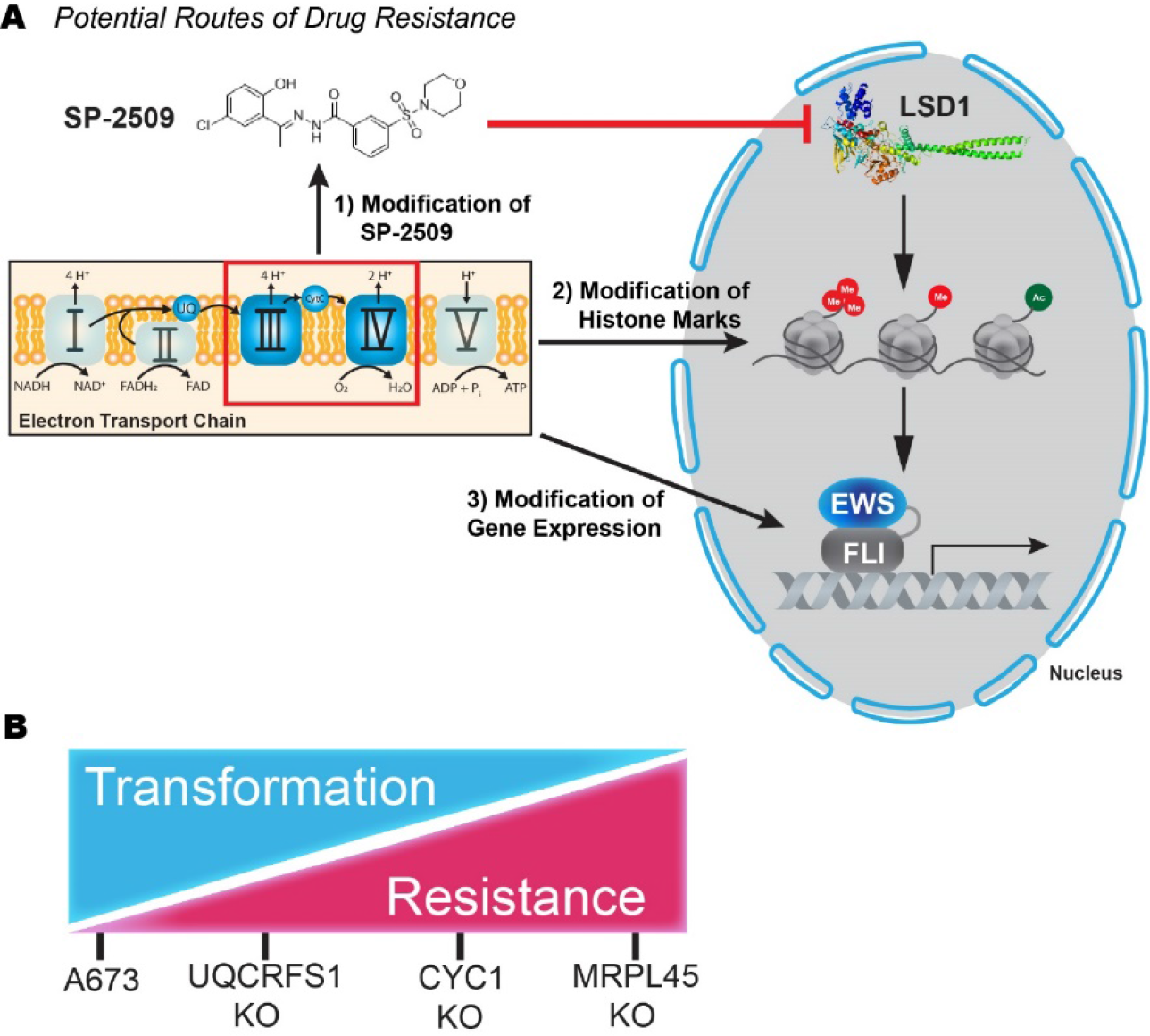
Model for mitochondrial influence on SP-2509 drug-resistance and oncogenic transformation ability in Ewing sarcoma. **A.** Each arrow indicates a potential route for the mitochondrial ETC to influence SP-2509 treatment of Ewing sarcoma. **B.** Transformation/SP-2509 drug-resistance model for mitochondrial dysfunction in Ewing sarcoma with A673 and each monoclonal cell line indicated.

Additionally, an unknown off-target specificity of SP-2509 that directly targets a component of mitochondrial function may exists that is unrelated to LSD1 inhibition. Though this is not an exhaustive list of potential drug resistance mechanisms, our work here allows us to address each of these potential pathways in future studies.

This study has defined mitochondrial dysfunction as a bona fide mechanism of drug resistance in Ewing sarcoma. Here, we have shown that loss of specific complexes of the mitochondria, particularly ETC CIII and CIV, causes resistance to SP-2509. However, these proteins are essential for proper cellular function, and we showed that disruption of mitochondrial protein function also alters oncogenic transformation (Supplementary Figure S7). Thus, a trade-off may occur in Ewing sarcoma cells that have mitochondrial dysfunction, whereby they gain the ability to resist SP-2509/seclidemstat treatment but lose their ability to survive and proliferate in a tumor microenvironment (Figure 5B). This spectrum of transformation/drug resistance is exemplified by our monoclonal KO cells with our most resistant cell line MRPL45 KO, unable to undergo oncogenic transformation, whereas our least resistant cell line UQCRFS1 KO retains the ability to undergo oncogenic transformation. CYC1 KO cells are moderately resistant to SP-

2509, and still retain some ability to undergo oncogenic transformation. Therefore, these data offer valuable insights into potential combination therapeutics that may promote dependence on mitochondrial metabolism, in particular oxidative phosphorylation, to promote more long-term efficacy of LSD1 inhibitors.

## Materials and Methods

### Cell lines and reagents

Cell lines were sourced and cultured as described previously (Johnson et al., 2017, Pishas et al., 2018, Theisen, Miller et al., 2019). Monoclonal cell lines generated in this study were expanded and stored long-term in 90% Fetal Bovine Serum (FBS) and 10% DMSO in liquid nitrogen. Soft agar assays were performed as described (Johnson et al., 2017, Pishas et al., 2018, Theisen et al., 2019). SP-2509 was purchased from Cayman Chemicals. Antimycin A, rotenone, and oligomycin were purchased from Thermo-Fisher. All oligonucleotides used in this study were purchased from Integrate DNA Technologies (IDT), or Thermo-Fisher Scientific.

### Genome-scale CRISPR-Cas9 KO screen

A genome-scale CRISPR-Cas9 KO screen was performed (Dana-Farber Cancer Institute) using an Avana-4 lentiviral library with barcode sequences for sgRNAs located in Supplementary File 1. Ewing sarcoma cell lines, A673 and TC-32, cells were seeded and infected with the Avana-4 lentiviral library in 12-well tissue culture treated plates overnight before transfer to T-75 flasks. Cells were selected with puromycin and blasticidin for 6-7 days to ensure stable expression of Cas9 and sgRNA, respectively. Selection media was removed, and the cells recovered for 24 hr before treatment with either DMSO or 300 nM SP-2509 (n ≥ 2 for each condition). An early time point for each condition was collected, and the remaining cells were passaged for 2 weeks before the final collection. Genomic DNA was isolated, PCR amplified, and subjected to Illumina sequencing. The 5′ and 3′ PCR primers included the P5 (5’-AATGATACGGCGACCACCGA-3’) and P7 (5’-CAAGCAGAAGACGGCATACGAGAT-3’) Illumina adapter sequences. PoolQ (Broad Institute) was used to deconvolute samples by barcode and quantify the results.

### Cloning, and lentivirus production

All constructs used to target genes with CRISPR-Cas9 were produced by following the ‘Lentiviral CRISPR Toolbox’ by Feng Zhang (Shalem, Sanjana et al., 2014), with some minor modifications. To generate CRISPR-Cas9 constructs, a Control_sg plasmid containing the sgRNA under control of a U6 promoter and spCas9-NLS-1XFLAG-P2A-puro under the control of an EFS-NS promoter was used. Control_sg was digested with *BsmB1* and dephosphorylated overnight. The linear constructs were purified via agarose gel electrophoresis and QIAquick Gel Extraction Kit (Qiagen). Insert oligonucleotides were designed based on Avana-4 library single guide RNA (sgRNA) sequences (Supplementary Table S3 and Supplementary File 1) (Shalem et al., 2014). Insert oligonucleotides were phosphorylated using T4 polynucleotide kinase (NEB) at 37 °C for 1 hr and were annealed by heating to 95°C for 5 min and slowly cooled to room temperature. Annealed and phosphorylated oligonucleotides were diluted (1:200) and ligation reactions were performed at room temperature overnight and transformed into DH5α chemically competent *Escherichia coli*. Individual colonies were grown overnight, purified by Qiagen miniprep, and Sanger sequenced (Eurofins Scientific) to confirm insertion of desired sgRNA. Lentivirus was produced by transfecting EBNA (HEK-293) cells with 10 µg each of a CRISPR- Cas9 construct, LentiVSVG, and psPAX2 in OptiMEM and *Trans*IT-LT1 (Mirus). Transfected EBNA cells were incubated at 37 °C for 48 hr, and virus was collected in modified DMEM. Virus was filtered (2 µm) and stored at -80 °C.

### CRISPR-Cas9 lentivirus transduction

A673 cells were seeded in 10 cm dishes (2x10^6^ cells) and allowed to adhere overnight at 37 °C. Cells were transduced by removing media and adding 2 mL of lentivirus and polybrene (10 µg/mL). Transduced plates were incubated at 37 °C with gentle rocking every 30 min. After 2 hr, media (6 mL) containing polybrene (10 µg/mL) was added and the cells were left for 48 hr. Cells were selected with puromycin (2 µg/mL) were incubated for at least 48 hr, or until all cells on a control (non-transduced) plate were dead.

### Generation of monoclonal cell lines

We generated long-term stable knockouts of mitochondrial genes by performing lentiviral transduction in A673 cells (see above) and performing limited dilution in 96-well plates. After transduction and 2 days of selection, cells were serially diluted in A673 media containing puromycin. Cells were incubated at 37 °C undisturbed for 1 week in 96 well plates. As single colonies reached confluency, they were transferred first to a 24-well plate, and eventually to a 10-cm dish. These monoclonal cell lines were assayed via western blot to determine efficiency of gene KO, and subsequently assayed for cell proliferation and cell viability to assess drug resistance phenotype. Frozen aliquots of monoclonal populations were stored in 90% FBS and 10% DMSO and frozen in liquid nitrogen.

### Western blot analysis

Antibodies for western blot analysis were sourced from either Thermo-Fisher (MRPL45, PA5- 54778), Abcam (COA4, ab105678; UQCRFS1, ab14746; CYC1, ab137757; NDUFS1, ab169540; ATP5F1, ab84625; SDHC, ab129736; Membrane Integrity WB Antibody Cocktail, ab110414; FLI (EWS/FLI), ab15289), or Cell Signaling Technology (LSD1, C69G12). Dilutions of antibodies were performed based on manufacturer’s protocol. Confirmation of gene knockout was performed as previously described (Pishas et al., 2018).

### Seahorse XF assays

Metabolic flux analyses were performed using a Seahorse XFp flux analyzer (Agilent) following manufacturer protocols. Optimal cell seeding densities were determined for each cell line and condition. For mitochondrial stress tests, inhibitors (oligomycin, FCCP, rotenone/antimycin A) were prepared according to manufacturer’s protocol (Agilent). Data were analyzed using Agilent Wave software and GraphPad Prism.

### Cell confluency using Incucyte live-cell analysis instrument

Cells were harvested and seeded in 96-well clear flat bottom tissue culture treated plates (Corning). Cells were seeded at 5,000-8,000 cells per well in 200 µL of media and adhered for at least 2 hr at 37 °C. Plates were analyzed in an Incucyte Live-Cell analysis instrument (Essen Biosciences) and phase contrast images were collected every 3 hr.

### RNA-sequencing analysis

Cells for RNA-sequencing (RNA-Seq) were prepared by seeding cells (1x10^6^) on a 10 cm tissue culture treated plate and were incubated overnight at 37 °C in 5% CO2. Media was removed and replaced with media containing either 0.1% DMSO or 500 nM SP-2509, each cell line and treatment condition had 2 replicates. After 48 hr of treatment with either DMSO or SP-2509, cells were harvested, and total mRNA was purified using the RNeasy kit (Qiagen) following manufacturer’s protocol, and on-column DNase digestion was performed. RNA was eluted from spin-columns using DEP-C treated dH2O, and RNA concentration was measured using QIAxpert (Qiagen). Duplicate RNA samples were submitted to the Institute for Genomic Medicine (IGM) (Abigail Wexner Research Institute at Nationwide Children’s Hospital, Columbus, OH 43205) following IGM sample submission guidelines. Samples were sequenced using a Hi-Seq 4000 to generate 150 bp paired-end reads. STAR (2.7) was used to align reads to the human genome build hg19 and generate read counts for each gene. Genes with < 1 count per sample were excluded from further analysis. DESeq2 was used for differential expression of gene counts. Volcano plots, principal component analysis, and heatmaps were generated in R. Heatmaps were generated using *pheatmap* with unsupervised hierarchical clustering. Scatterplots were generated using *ggplot2* and Pearson correlation coefficients were determined. Datasets were compared using Gene set enrichment analysis (GSEA). Pathway comparison analysis was performed using compiled gene sets from our RNA-seq data and those accessible from MsigDB.

## Data Availability

The datasets and computer code produced in this study are available in the following databases: RNA-sequencing raw and processed data are available under the GEO accession: GSE185379 Analytical pipelines: Available upon reasonable request.

## Acknowledgements

We would like to thank Ariunaa Bayanjargal, Megann Boone, Andrea Byrum, Julia Selich-Anderson, and Iftekhar Showpnil for helpful discussion and careful editing of this manuscript.

## Author Contributions

LMG, KIP, KS, and SLL are responsible for conceptualization of the project. Investigation was performed by EJT, JCC, JS, GR, BSJ, AK, and LMG, JS. Methodology was formulated by EJT, JCC, LMG, JS, KS, ERT and SLL. Data analysis was performed by EJT, JCC, LMG, JS, GR, BJ, AK, CT and GA. Software and data curation were performed by ERT, CT, and GA. Writing of the original draft was completed by EJT. Reviewing and editing was performed by all authors. Funding acquisition was completed by KIP, KS, SLL, and ERT. Supervision was provided by ERT and SLL.

## Abbreviations

Ewing sarcoma breakpoint region 1 protein, EWSR1; Friend leukemia integration 1 transcription factor, FLI1; BRG1-BRM-associated factor, BAF; nucleosome remodeling and deacetylase, NuRD; Lysine-specific demethylase-1, LSD1; REST corepressor, CoREST; electron transport chain, ETC; RNA-sequencing, RNA-seq; Fetal Bovine Serum, FBS; single guide RNA, sgRNA; gene set enrichment analysis, GSEA; log2 fold change, log2FC; open reading frames, ORFs; mitochondrial ribosomal protein, MRP; Mitochondrial Ribosome Protein L45, MRPL45; Ubiquinol-Cytochrome C Reductase, Rieske Iron-Sulfur Polypeptide 1, UQCRFS1; cytochrome C1, CYC1; Cytochrome C Oxidase Assembly Factor 4 Homolog, COA4; NADH:Ubiquinone Oxidoreductase Core Subunit S1, NDUFS1, succinate dehydrogenase c, SDHC, ATP synthase F(0) complex subunit B1, ATP5F1; YFP-tagged Parkin, YFP-Parkin; Inner-membrane space, IMS; Inner-membrane, IM; Normalized Enrichment Score, NES; gene set, GS; unfolded protein response; UPR; endoplasmic reticulum, ER; apoptotic protease activating factor-1, Apaf-1; gastrointestinal stromal tumors, GIST; Ten-eleven translocation dioxygenase, TET; α- ketoglutarate, α-KG; S-adenosyl methionine, SAM; Jumonji C domain-containing, JmjC

## Supplementary Data

### Supplementary Tables

**Supplementary Table S1:** Common genes identified in CRISPR-Cas9 screen for A673 and TC-32 cell lines.

| # | Core gene hit | # Guide hits A673 | # Guide hits TC-32 | # | Core gene hit | # Guide hits A673 | # Guide hits TC-32 |
| --- | --- | --- | --- | --- | --- | --- | --- |
| 1 | ALAD | 1 | 1 | 22 | LRPPRC | 3 | 2 |
| 2 | APOL6 | 1 | 1 | 23 | MRPL34 | 4 | 1 |
| 3 | ASF1A | 1 | 1 | 24 | NEUROG1 | 1 | 1 |
| 4 | ATXN7 | 1 | 2 | 25 | NPS | 1 | 1 |
| 5 | BAX | 1 | 1 | 26 | PTPN18 | 1 | 1 |
| 6 | BOLA3 | 3 | 1 | 27 | SCG2 | 1 | 1 |
| 7 | COQ7 | 4 | 1 | 28 | SCO2 | 3 | 1 |
| 8 | COX4I1 | 4 | 1 | 29 | SEMG2 | 1 | 1 |
| 9 | COX8A | 3 | 1 | 30 | SERPINB4 | 1 | 1 |
| 10 | DOT1L | 2 | 1 | 31 | SIGLEC5 | 1 | 1 |
| 11 | E2F8 | 1 | 1 | 32 | SLC25A18 | 1 | 1 |
| 12 | EP300 | 1 | 1 | 33 | SLC30A3 | 1 | 1 |
| 13 | ERCC5 | 1 | 1 | 34 | SLFN11 | 1 | 1 |
| 14 | FAHD1 | 1 | 1 | 35 | STX3 | 1 | 1 |
| 15 | FRG2C | 1 | 1 | 36 | TAF6L | 2 | 1 |
| 16 | GLRX5 | 4 | 1 | 37 | TNP1 | 1 | 1 |
| 17 | HMBS | 4 | 1 | 38 | UQCC1 | 3 | 1 |
| 18 | IBA57 | 4 | 2 | 39 | UQCC2 | 2 | 1 |
| 19 | ISCA2 | 4 | 3 | 40 | YME1L1 | 1 | 1 |
| 20 | KLHL40 | 1 | 1 | 41 | ZRANB3 | 1 | 1 |
| 21 | L3MBTL3 | 1 | 2 |  |  |  |  |

**Supplementary Table S2:**
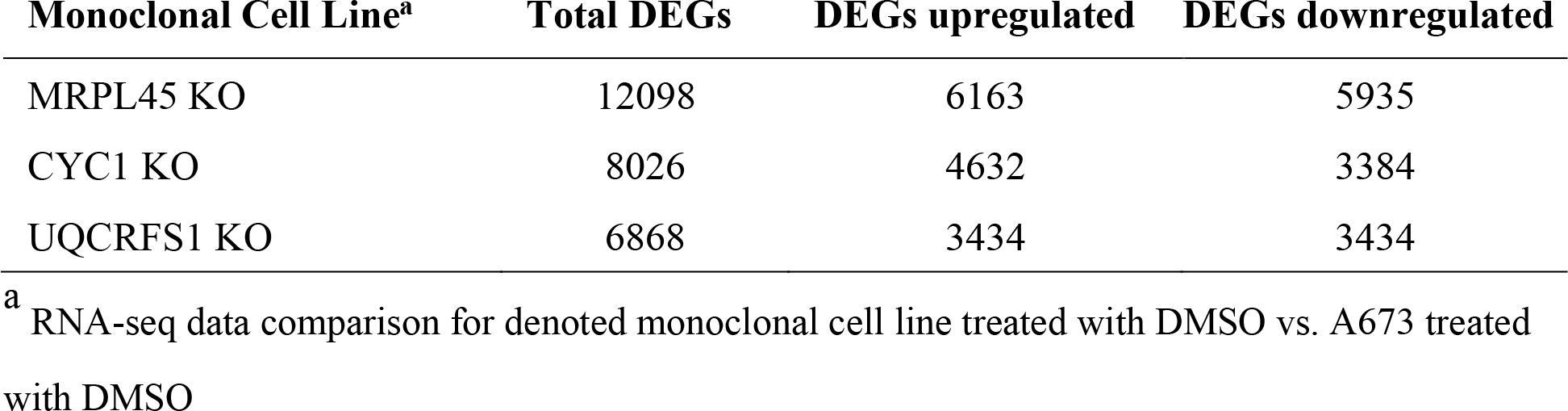
Basal transcriptional changes induced by KO in monoclonal cell lines.

**Supplementary Table S3:**
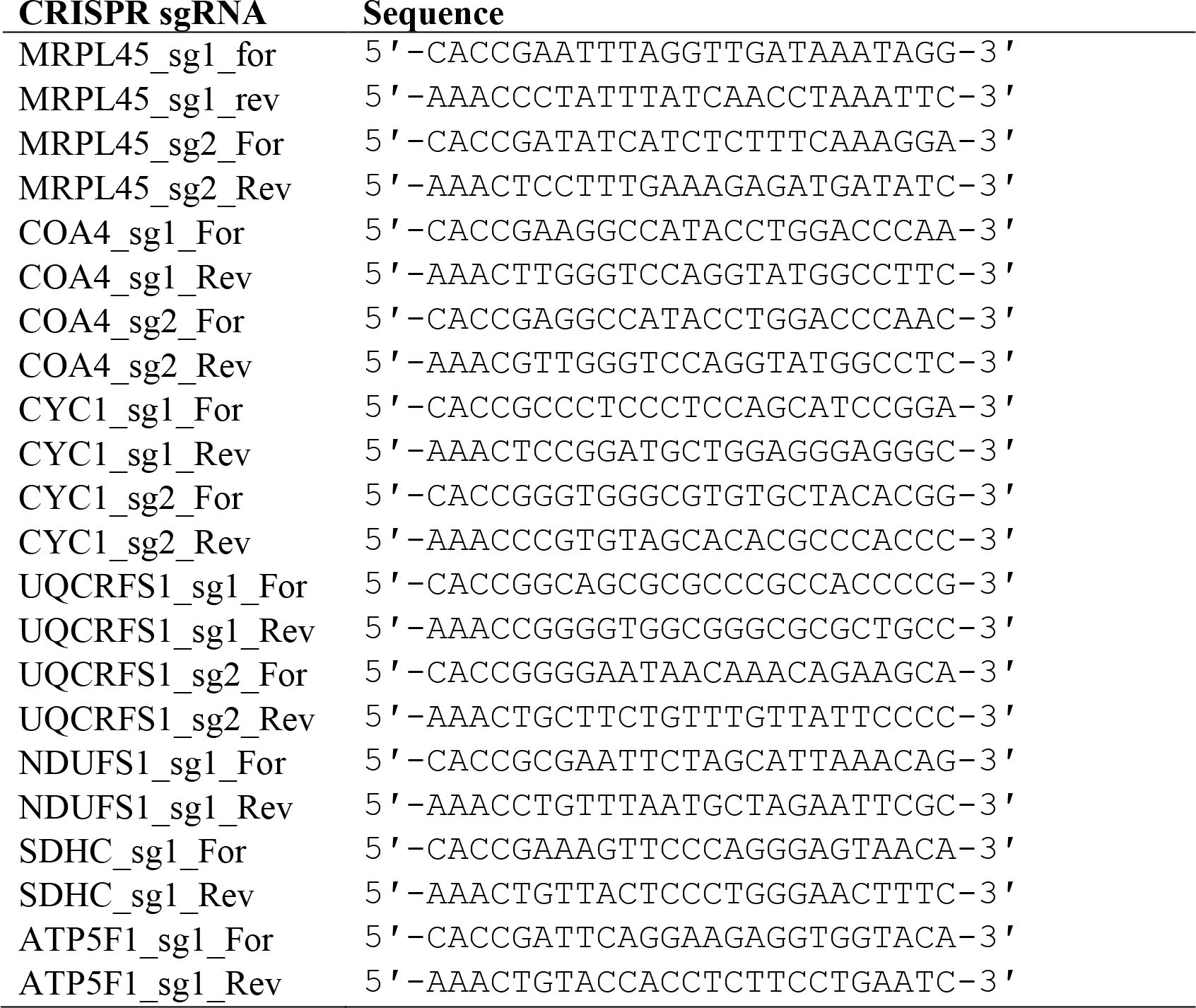
DNA oligo inserts used to generate indicated CRISPR-Cas9 gene KO.

### Supplementary Figure Legends

**Supplementary Figure S1.**
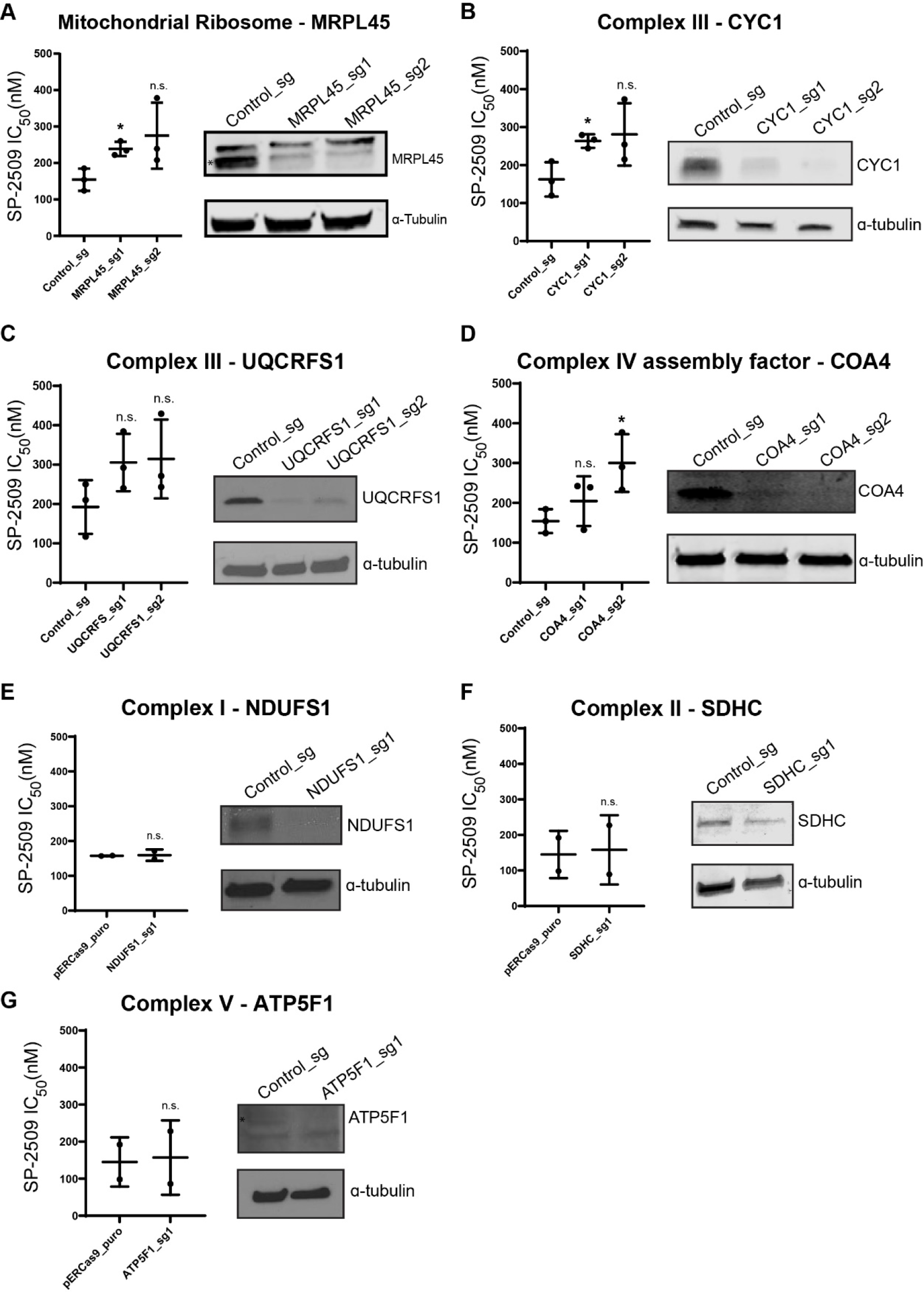
Validation of CRISPR screen with KO of various mitochondrial genes. Compiled cell viability assays and western blots for A673 cells transduced with lentivirus containing constructs for KO of *MRPL45* (**A)**, *CYC1* (**B**), *UQCRFS1* (**C**), *COA4* (**D**), *NDUFS1* (**E**), *SDHC* (**F**), and *ATP5F1* (**G**) compared to control infection, Control_sg. Cells were treated with SP-2509 (0.01 – 30 µM) for 72 hr. Data represent mean ± standard deviation from three technical replicates. Statistical analysis was performed with a student’s t-test with a * indicating a p-value ≤ 0.05, and values that failed to reach significance are marked not significant (n.s.). For blots with multiple protein bands, the band marked with a * indicates the proper size of the target protein.

**Supplementary Figure S2.**
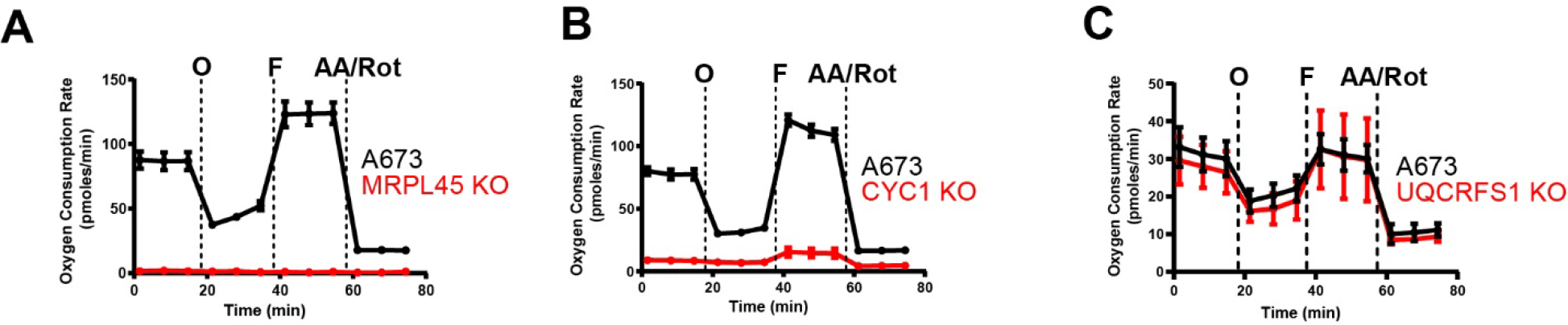
Oxidative phosphorylation capacity of monoclonal KO cells. Seahorse XF mitochondrial stress tests comparing A673 cells with MRPL45 KO (**A**), CYC1 KO (**B**), and UQCRFS1 KO **(C)**. Spike-in of oligomycin, FCCP, and a combination of antimycin A and rotenone occur as indicated.

**Supplementary Figure S3.**
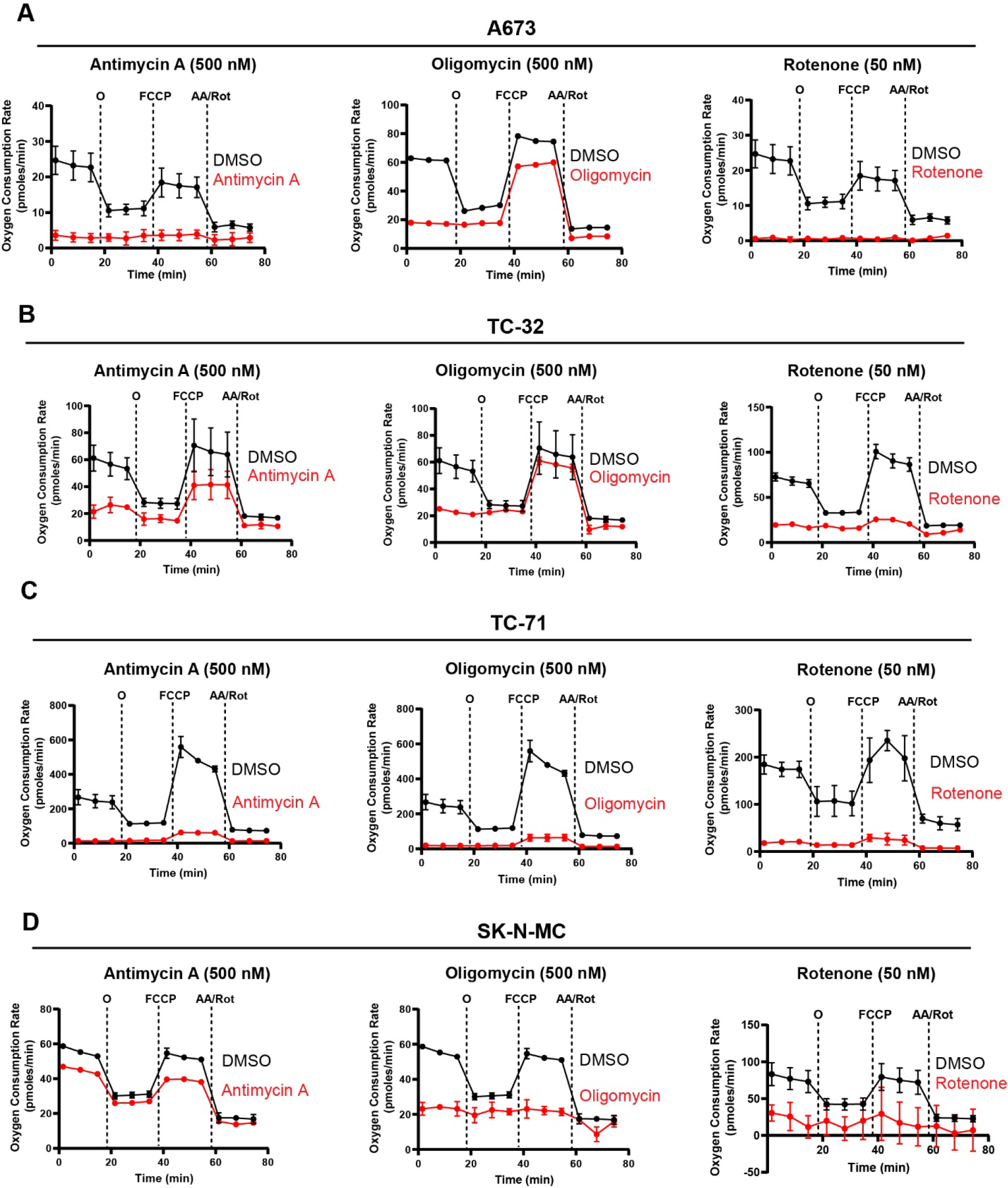
Seahorse analysis of Ewing sarcoma cell lines in the presence of ETC inhibitors. Seahorse XF mitochondrial stress tests performed on A673 (**A**), TC-32 (**B**), TC-71 (**C**), and SK- N-MC (**D**). Each cell line was treated with the either antimycin A (500 nM), oligomycin (500 nM) or rotenone (50 nM) for 72 hrs. Spike-in of oligomycin, FCCP, and a combination of antimycin A and rotenone occur as indicated.

**Supplementary Figure S4.**
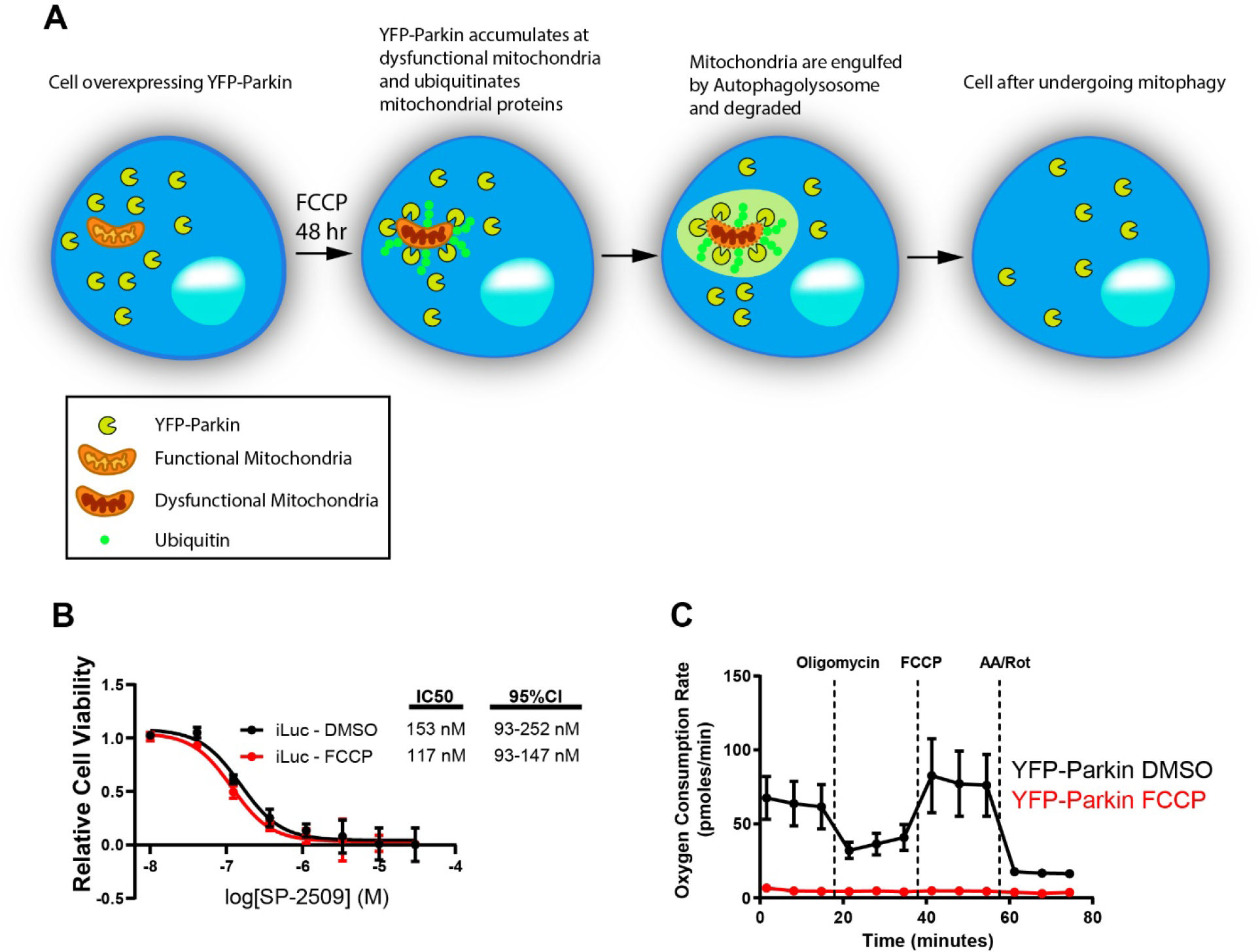
Depletion of mitochondria from A673 cells. **A.** A673 cells stably expressing YFP-Parkin are treated with the proton uncoupler, Trifluoromethoxy carbonylcyanide phenylhydrazone (FCCP), for 48 hr, which induces mitochondrial stress. Parkin accumulates on the outer membrane of dysfunctional mitochondria leading to the ubiquitination of mitochondrial proteins. This signals the cell to direct the dysfunctional mitochondria to the lysosome through mitophagy. **B.** Dose response curves for A673 cells transduced with a mock retroviral infection (iLuc), treated with either DMSO (black) or FCCP (red). Cells were treated with SP-2509 (0.01 – 30 µM) for 72 hr. Data represent mean ± standard deviation from three replicates. **C.** Mitochondrial stress tests using a Seahorse XFp comparing A673 cells with mitochondrial null (mtø). Three measurements were taken initially to measure the basal oxygen consumption rate, and three measurements each were taken following spike-in of oligomycin, FCCP, and a combination of antimycin A and rotenone (as indicated).

**Supplementary Figure S5.**
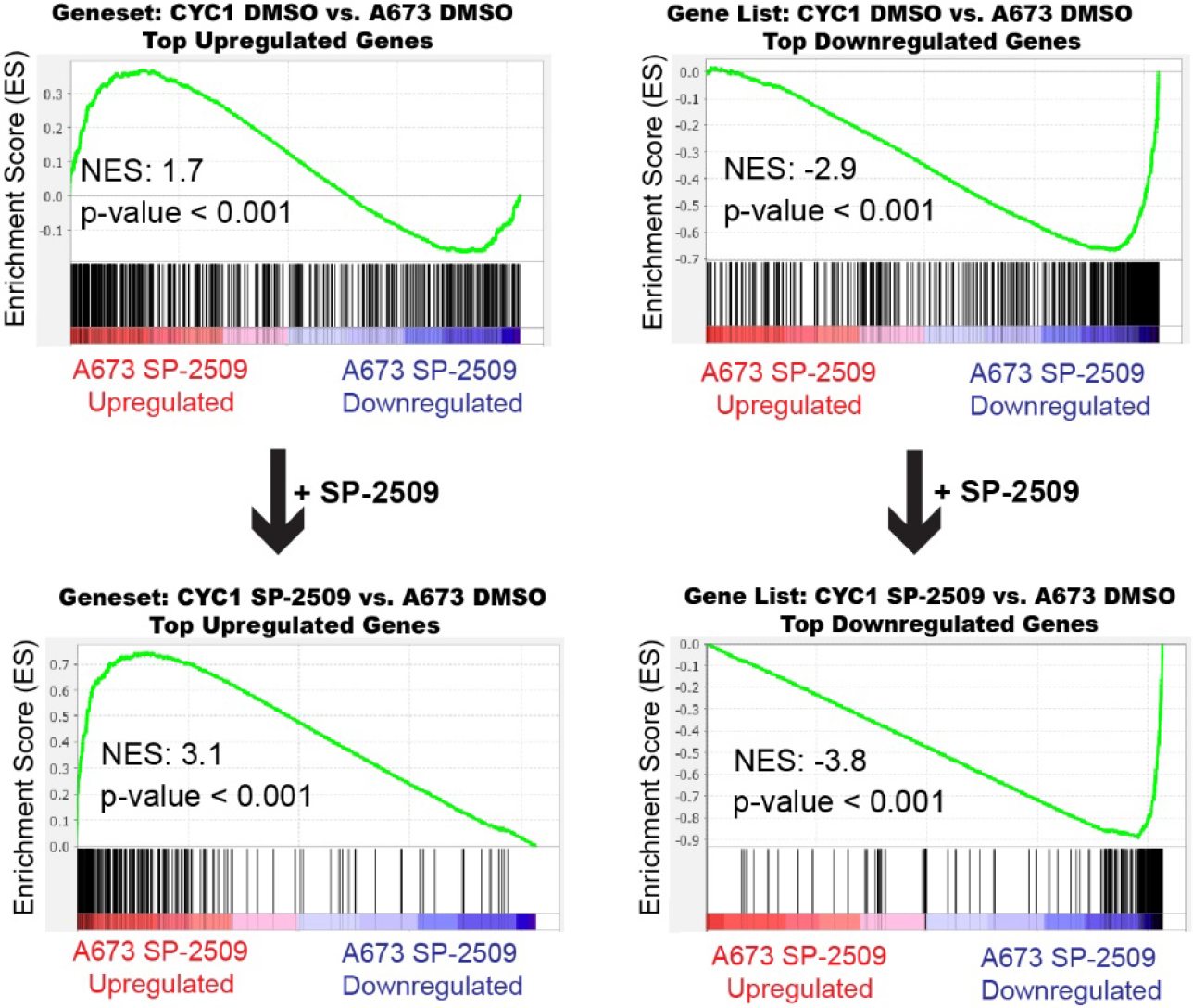
Gene Set Enrichment Analysis (GSEA) of monoclonal CYC1 KO compared to SP-2509 treated A673 cells. Gene set enrichment analysis (GSEA) from RNA-seq experiments using the LSD1 (SP-2509 treatment) regulated genes in A673 cells as the rank-ordered dataset for each comparison to each gene set for CYC1 KO. The top panel represents comparisons for the 480 most upregulated and downregulated genes for CYC1 KO treated with DMSO. The bottom panel represents GSEA for the most upregulated and downregulated genes for the indicated monoclonal KO treated with SP- 2509. Arrows indicate changes in gene expression from DMSO treated cells to SP-2509 treated cells. Normalized enrichment scores (NES) and p-values are shown for each GSEA.

**Supplementary Figure S6.**
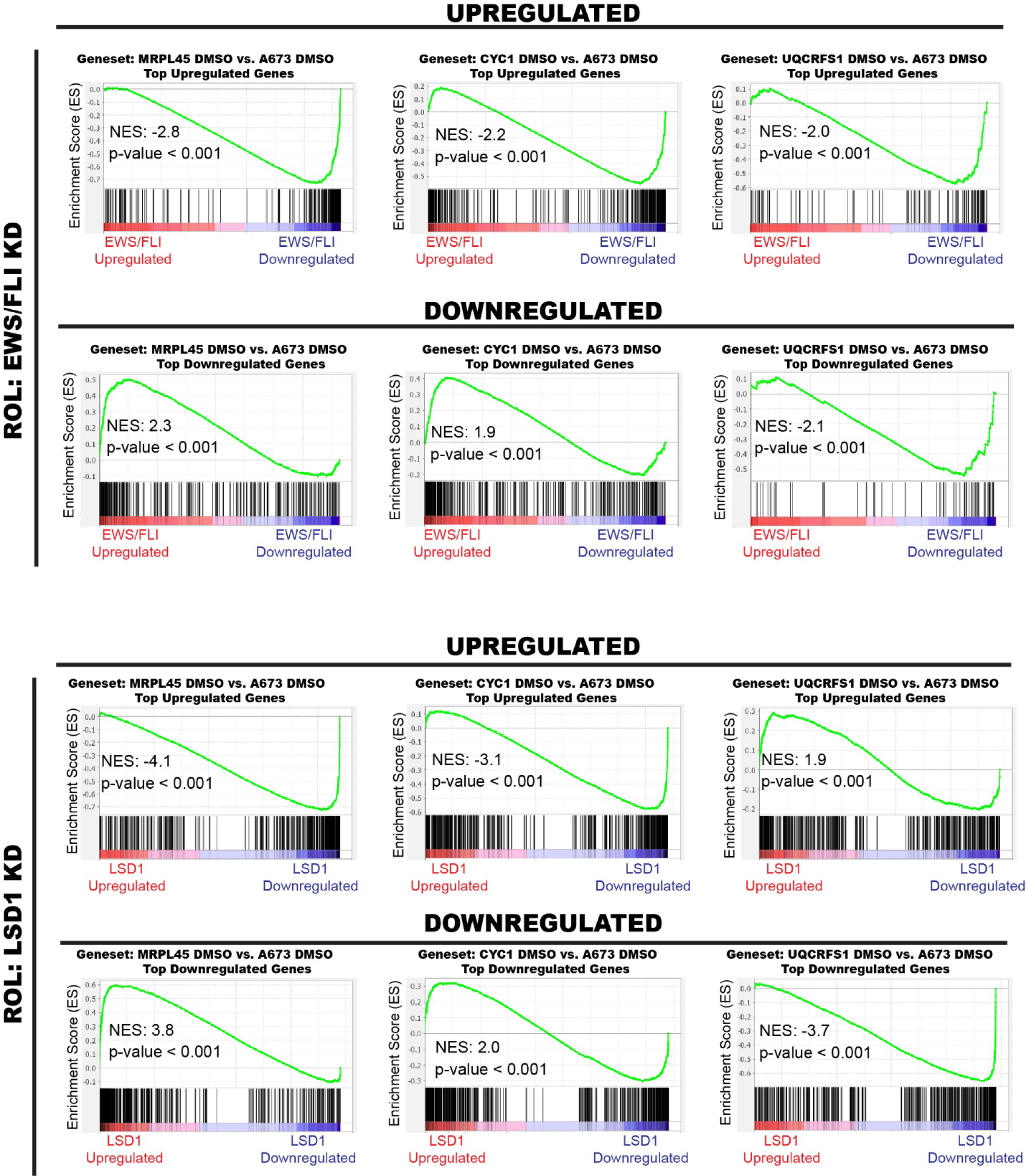
GSEA for monoclonal KO cells compared to EWS/FLI and LSD1 rank-ordered lists. Gene set enrichment analysis (GSEA) from RNA-seq experiments using the EWS/FLI or LSD1 regulated genes in A673 cells. The rank-ordered list for EWS/FLI (EWS/FLI knockdown - Sankar and Theisen et al. 2014) and LSD1 (LSD1 knockdown – Pishas et al. 2018) were compared to each gene set for MRPL45 KO, CYC1 KO, and UQCRFS1 KO as indicated. Each geneset represents the top 480 most upregulated or downregulated genes for the indicated monoclonal KO compared to A673 DMSO. Normalized Enrichment Scores (NES) and p-values are indicated for each comparison.

**Supplementary Figure S7.**
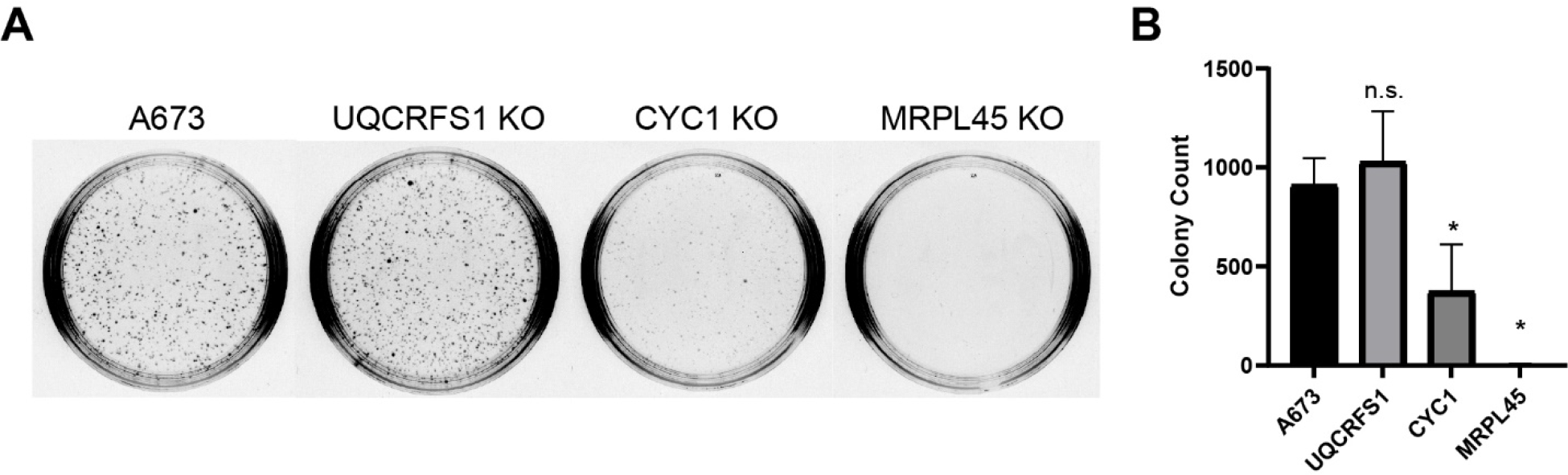
Oncogenic transformation potential of monoclonal KO cells. **A.** Colony formation assays for A673 and each indicated monoclonal cell line with images taken after 2 weeks post-seeding. **B.** Quantification for three independent colony formation assays. Statistical analysis was performed with a student’s t-test, with * denoting p-value ≤ 0.05 and values that are not significant denoted as n.s.

**Supplementary Figure S8.**
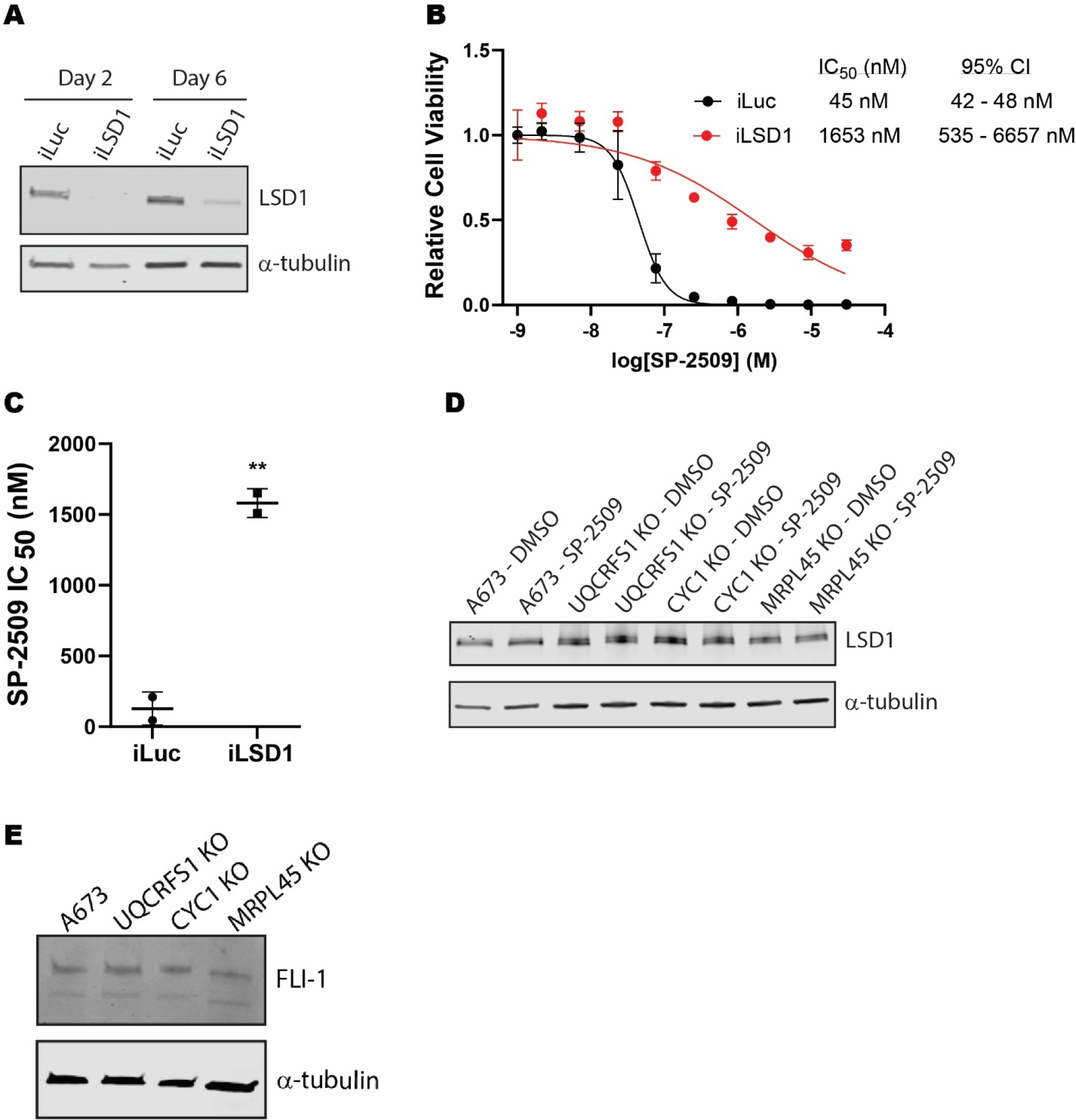
Knockdown of LSD1 confers resistance to SP-2509. **A.** Western blots for A673 cells transduced with retrovirus containing constructs for knockdown of *LSD1* (iLSD1) control infection, iLuc. **B.** Cell viability assay comparing iLSD1 to iLuc. Cells were treated with SP-2509 (0.01 – 30 µM) for 72 hr. Data represent mean ± standard deviation from three technical replicates. **C.** Compiled IC50 from cell viability assays comparing iLuc and iLSD1. Statistical analysis was performed with a student’s t-test with ** indicating a p-value ≤ 0.005. **D.** Western blot detecting expression levels of *LSD1* in monoclonal KO cells. Conditions are indicated for either 0.1% DMSO or 300 nM SP-2509 treatment (48 hr). **E.** Western blot detecting expression levels of *FLI1* (*EWS/FLI*) in monoclonal KO cells.

## References

1. Bennani-Baiti IM, Machado I, Llombart-Bosch A, Kovar H (2012) Lysine-specific demethylase 1 (LSD1/KDM1A/AOF2/BHC110) is expressed and is an epigenetic drug target in chondrosarcoma, Ewing’s sarcoma, osteosarcoma, and rhabdomyosarcoma. Hum Pathol 43: 1300–7

2. Boulay G, Sandoval GJ, Riggi N, Iyer S, Buisson R, Naigles B, Awad ME, Rengarajan S, Volorio A, McBride MJ, Broye LC, Zou L, Stamenkovic I, Kadoch C, Rivera MN (2017) Cancer-Specific Retargeting of BAF Complexes by a Prion-like Domain. Cell 171: 163–178 e19

3. Brohl AS, Solomon DA, Chang W, Wang J, Song Y, Sindiri S, Patidar R, Hurd L, Chen L, Shern JF, Liao H, Wen X, Gerard J, Kim JS, Lopez Guerrero JA, Machado I, Wai DH, Picci P, Triche T, Horvai AE et al. (2014) The genomic landscape of the Ewing Sarcoma family of tumors reveals recurrent STAG2 mutation. PLoS Genet 10: e1004475

4. Burningham Z, Hashibe M, Spector L, Schiffman JD (2012) The epidemiology of sarcoma. Clin Sarcoma Res 2: 14

5. Chakrabarty RP, Chandel NS (2021) Mitochondria as Signaling Organelles Control Mammalian Stem Cell Fate. Cell Stem Cell 28: 394–408

6. Chen Y, Yang Y, Wang F, Wan K, Yamane K, Zhang Y, Lei M (2006) Crystal structure of human histone lysine-specific demethylase 1 (LSD1). Proc Natl Acad Sci U S A 103: 13956–61

7. Correia-Melo C, Ichim G, Tait SW, Passos JF (2017) Depletion of mitochondria in mammalian cells through enforced mitophagy. Nat Protoc 12: 183–194

8. Crompton BD, Stewart C, Taylor-Weiner A, Alexe G, Kurek KC, Calicchio ML, Kiezun A, Carter SL, Shukla SA, Mehta SS, Thorner AR, de Torres C, Lavarino C, Sunol M, McKenna A, Sivachenko A, Cibulskis K, Lawrence MS, Stojanov P, Rosenberg M et al. (2014) The genomic landscape of pediatric Ewing sarcoma. Cancer Discov 4: 1326–41

9. Delattre O, Zucman J, Plougastel B, Desmaze C, Melot T, Peter M, Kovar H, Joubert I, de Jong P, Rouleau G, et al. (1992) Gene fusion with an ETS DNA-binding domain caused by chromosome translocation in human tumours. Nature 359: 162–5

10. Diebold LP, Gil HJ, Gao P, Martinez CA, Weinberg SE, Chandel NS (2019) Mitochondrial complex III is necessary for endothelial cell proliferation during angiogenesis. Nat Metab 1: 158–171

11. Doench JG, Fusi N, Sullender M, Hegde M, Vaimberg EW, Donovan KF, Smith I, Tothova Z, Wilen C, Orchard R, Virgin HW, Listgarten J, Root DE (2016) Optimized sgRNA design to maximize activity and minimize off-target effects of CRISPR-Cas9. Nat Biotechnol 34: 184–191

12. Gammage PA, Frezza C (2019) Mitochondrial DNA: the overlooked oncogenome? BMC Biol 17: 53

13. Gaspar N, Hawkins DS, Dirksen U, Lewis IJ, Ferrari S, Le Deley MC, Kovar H, Grimer R, Whelan J, Claude L, Delattre O, Paulussen M, Picci P, Sundby Hall K, van den Berg H, Ladenstein R, Michon J, Hjorth L, Judson I, Luksch R, et al. (2015) Ewing Sarcoma: Current Management and Future Approaches Through Collaboration. J Clin Oncol 33: 3036–46

14. Groenendijk FH, Bernards R (2014) Drug resistance to targeted therapies: deja vu all over again. Mol Oncol 8: 1067–83

15. Grunewald TGP, Cidre-Aranaz F, Surdez D, Tomazou EM, de Alava E, Kovar H, Sorensen PH, Delattre O, Dirksen U (2018) Ewing sarcoma. Nat Rev Dis Primers 4: 5

16. Johnson KM, Mahler NR, Saund RS, Theisen ER, Taslim C, Callender NW, Crow JC, Miller KR, Lessnick SL (2017) Role for the EWS domain of EWS/FLI in binding GGAA-microsatellites required for Ewing sarcoma anchorage independent growth. Proc Natl Acad Sci U S A 114: 9870–9875

17. Killian JK, Kim SY, Miettinen M, Smith C, Merino M, Tsokos M, Quezado M, Smith WI, Jr., Jahromi MS, Xekouki P, Szarek E, Walker RL, Lasota J, Raffeld M, Klotzle B, Wang Z, Jones L, Zhu Y, Wang Y, Waterfall JJ et al. (2013) Succinate dehydrogenase mutation underlies global epigenomic divergence in gastrointestinal stromal tumor. Cancer Discov 3: 648–57

18. Lawrence MS, Stojanov P, Polak P, Kryukov GV, Cibulskis K, Sivachenko A, Carter SL, Stewart C, Mermel CH, Roberts SA, Kiezun A, Hammerman PS, McKenna A, Drier Y, Zou L, Ramos AH, Pugh TJ, Stransky N, Helman E, Kim J et al. (2013) Mutational heterogeneity in cancer and the search for new cancer-associated genes. Nature 499: 214–218

19. Liu Z, Butow RA (2006) Mitochondrial retrograde signaling. Annu Rev Genet 40: 159–85

20. Martinez-Reyes I, Chandel NS (2020) Mitochondrial TCA cycle metabolites control physiology and disease. Nat Commun 11: 102

21. Mason EF, Hornick JL (2013) Succinate dehydrogenase deficiency is associated with decreased 5-hydroxymethylcytosine production in gastrointestinal stromal tumors: implications for mechanisms of tumorigenesis. Mod Pathol 26: 1492–7

22. Meyers RM, Bryan JG, McFarland JM, Weir BA, Sizemore AE, Xu H, Dharia NV, Montgomery PG, Cowley GS, Pantel S, Goodale A, Lee Y, Ali LD, Jiang G, Lubonja R, Harrington WF, Strickland M, Wu T, Hawes DC, Zhivich VA et al. (2017) Computational correction of copy number effect improves specificity of CRISPR-Cas9 essentiality screens in cancer cells. Nat Genet 49: 1779–1784

23. Pishas KI, Drenberg CD, Taslim C, Theisen ER, Johnson KM, Saund RS, Pop IL, Crompton BD, Lawlor ER, Tirode F, Mora J, Delattre O, Beckerle MC, Callen DF, Sharma S, Lessnick SL (2018) Therapeutic Targeting of KDM1A/LSD1 in Ewing Sarcoma with SP-2509 Engages the Endoplasmic Reticulum Stress Response. Mol Cancer Ther 17: 1902–1916

24. Pishas KI, Lessnick SL (2018) Ewing sarcoma resistance to SP-2509 is not mediated through KDM1A/LSD1 mutation. Oncotarget 9: 36413–36429

25. Riggi N, Knoechel B, Gillespie SM, Rheinbay E, Boulay G, Suva ML, Rossetti NE, Boonseng WE, Oksuz O, Cook EB, Formey A, Patel A, Gymrek M, Thapar V, Deshpande V, Ting DT, Hornicek FJ, Nielsen GP, Stamenkovic I, Aryee MJ et al. (2014) EWS-FLI1 utilizes divergent chromatin remodeling mechanisms to directly activate or repress enhancer elements in Ewing sarcoma. Cancer Cell 26: 668–681

26. Sankar S, Bell R, Stephens B, Zhuo R, Sharma S, Bearss DJ, Lessnick SL (2013) Mechanism and relevance of EWS/FLI-mediated transcriptional repression in Ewing sarcoma. Oncogene 32: 5089–100

27. Sankar S, Theisen ER, Bearss J, Mulvihill T, Hoffman LM, Sorna V, Beckerle MC, Sharma S, Lessnick SL (2014) Reversible LSD1 inhibition interferes with global EWS/ETS transcriptional activity and impedes Ewing sarcoma tumor growth. Clin Cancer Res 20: 4584–97

28. Sarmento-Ribeiro AB, Scorilas A, Goncalves AC, Efferth T, Trougakos IP (2019) The emergence of drug resistance to targeted cancer therapies: Clinical evidence. Drug Resist Updat 47: 100646

29. Shalem O, Sanjana NE, Hartenian E, Shi X, Scott DA, Mikkelson T, Heckl D, Ebert BL, Root DE, Doench JG, Zhang F (2014) Genome-scale CRISPR-Cas9 knockout screening in human cells. Science 343: 84–87

30. Shi Y, Lan F, Matson C, Mulligan P, Whetstine JR, Cole PA, Casero RA, Shi Y (2004) Histone demethylation mediated by the nuclear amine oxidase homolog LSD1. Cell 119: 941–53

31. Sorna V, Theisen ER, Stephens B, Warner SL, Bearss DJ, Vankayalapati H, Sharma S (2013) High-throughput virtual screening identifies novel N’-(1-phenylethylidene)-benzohydrazides as potent, specific, and reversible LSD1 inhibitors. J Med Chem 56: 9496–508

32. Subramanian A, Tamayo P, Mootha VK, Mukherjee S, Ebert BL, Gillette MA, Paulovich A, Pomeroy SL, Golub TR, Lander ES, Mesirov JP (2005) Gene set enrichment analysis: a knowledge-based approach for interpreting genome-wide expression profiles. Proc Natl Acad Sci U S A 102: 15545–50

33. Sulahian R, Kwon JJ, Walsh KH, Pailler E, Bosse TL, Thaker M, Almanza D, Dempster JM, Pan J, Piccioni F, Dumont N, Gonzalez A, Rennhack J, Nabet B, Bachman JA, Goodale A, Lee Y, Bagul M, Liao R, Navarro A et al. (2019) Synthetic Lethal Interaction of SHOC2 Depletion with MEK Inhibition in RAS-Driven Cancers. Cell Rep 29: 118–134 e8

34. Szymczyna BR, Arrowsmith CH (2000) DNA binding specificity studies of four ETS proteins support an indirect read-out mechanism of protein-DNA recognition. J Biol Chem 275: 28363–70

35. Taanman JW (1999) The mitochondrial genome: structure, transcription, translation and replication. Biochim Biophys Acta 1410: 103–23

36. Theisen ER, Miller KR, Showpnil IA, Taslim C, Pishas KI, Lessnick SL (2019) Transcriptomic analysis functionally maps the intrinsically disordered domain of EWS/FLI and reveals novel transcriptional dependencies for oncogenesis. Genes Cancer 10: 21–38

37. Theisen ER, Pishas KI, Saund RS, Lessnick SL (2016) Therapeutic opportunities in Ewing sarcoma: EWS-FLI inhibition via LSD1 targeting. Oncotarget 7: 17616–30

38. Theisen ER, Selich-Anderson J, Miller KR, Tanner JM, Taslim C, Pishas KI, Sharma S, Lessnick SL (2021) Chromatin profiling reveals relocalization of lysine-specific demethylase 1 by an oncogenic fusion protein. Epigenetics 16: 405–424

39. Turc-Carel C, Aurias A, Mugneret F, Lizard S, Sidaner I, Volk C, Thiery JP, Olschwang S, Philip I, Berger MP, et al. (1988) Chromosomes in Ewing’s sarcoma. I. An evaluation of 85 cases of remarkable consistency of t(11;22)(q24;q12). Cancer Genet Cytogenet 32: 229–38

40. Turc-Carel C, Philip I, Berger MP, Philip T, Lenoir GM (1984) Chromosome study of Ewing’s sarcoma (ES) cell lines. Consistency of a reciprocal translocation t(11;22)(q24;q12). Cancer Genet Cytogenet 12: 1–19

41. Wang Y, Zhang H, Chen Y, Sun Y, Yang F, Yu W, Liang J, Sun L, Yang X, Shi L, Li R, Li Y, Zhang Y, Li Q, Yi X, Shang Y (2009) LSD1 is a subunit of the NuRD complex and targets the metastasis programs in breast cancer. Cell 138: 660–72

42. Wei GH, Badis G, Berger MF, Kivioja T, Palin K, Enge M, Bonke M, Jolma A, Varjosalo M, Gehrke AR, Yan J, Talukder S, Turunen M, Taipale M, Stunnenberg HG, Ukkonen E, Hughes TR, Bulyk ML, Taipale J (2010) Genome-wide analysis of ETS-family DNA-binding in vitro and in vivo. EMBO J 29: 2147–60

43. Weinberg SE, Singer BD, Steinert EM, Martinez CA, Mehta MM, Martinez-Reyes I, Gao P, Helmin KA, Abdala-Valencia H, Sena LA, Schumacker PT, Turka LA, Chandel NS (2019) Mitochondrial complex III is essential for suppressive function of regulatory T cells. Nature 565: 495–499

44. Yang D, Kim J (2019) Mitochondrial Retrograde Signalling and Metabolic Alterations in the Tumour Microenvironment. Cells 8

